# Remote ischemic conditioning attenuates transneuronal degeneration and promotes stroke recovery via CD36-mediated efferocytosis

**DOI:** 10.1101/2024.09.03.611124

**Authors:** Hyunwoo Ju, Il-doo Kim, Ina Pavlova, Shang Mu, Keun Woo Park, Joseph Minkler, Ahmed Madkoor, Wei Wang, Xiaoman Wang, Zhuhao Wu, Jiwon Yang, Maria Febbraio, John W. Cave, Sunghee Cho

**Affiliations:** Burke Neurological Institute, 785 Mamaroneck Ave, White Plains, NY 10605, USA; Helen & Robert Appel Alzheimer’s Disease Research Institute, Weill Cornell Medicine, 413 E 69th St, New York, NY 10021, USA; Department of Dentistry and Dental Hygiene, University of Alberta, Edmonton, Alberta, Canada; InVitro Cell Research, Englewood, NJ, USA; Feil Brain Mind Research Institute, Weill Cornell Medicine, 1600 York Avenue, New York, NY, USA

**Keywords:** Ischemic stroke, Injury resolution, Monocyte/Macrophages, Clearance, Cellular debris

## Abstract

**BACKGROUND:** Remote ischemic limb conditioning (RIC) has been implicated in cross-organ protection in cerebrovascular disease, including stroke. However, the lack of a consensus protocol and controversy over the clinical therapeutic outcomes of RIC suggest inadequate mechanistic understanding of RIC. The current study identifies RIC-induced molecular and cellular events in the blood that enhance long-term functional recovery in experimental cerebral ischemia

**METHODS:** Naive mice or mice subjected to transient ischemic stroke were randomly selected to receive sham conditioning or RIC in the hind limb at 2 h post-stroke. At 3d post-stroke, monocyte composition in the blood was analyzed, and brain tissue was examined for monocyte-derived macrophages (Mφ), levels of efferocytosis, and CD36 expression. Mouse with conditional deletion of CD36 in Mφ (cKO^MMφ^) was used to establish the role of CD36 in RIC-mediated modulation of efferocytosis, transneuronal degeneration, and recovery following stroke.

**RESULTS:** RIC applied 2h after stroke increased entry of monocytes into the injured brain. In the post-ischemic brain, Mφ had increased levels of CD36 expression and efferocytosis. These changes in brain Mφ were derived from RIC-induced changes in circulating monocytes. In the blood, RIC increased CD36 expression in circulating monocytes and shifted monocytes to a proinflammatory LY6C^High^ state. Conditional deletion of CD36 in Mφ abrogated the RIC-induced monocyte shift in the blood and efferocytosis in the brain. During the recovery phase of stroke, RIC rescued the loss of the volume and of tyrosine hydroxylase+ neurons in substantia nigra (SN) as well as behavioral deficits in WT mice, but not in cKO^MMφ^ mice.

**CONCLUSIONS:** RIC induces a shift in monocytes to a proinflammatory state with elevated CD36 levels, and this is associated with CD36-dependent efferocytosis in Mφs that rescues delayed transneuronal degeneration in the post-ischemic brain and promotes stroke recovery. Together, these findings provide novel insight into our mechanistic understanding of how RIC improves in post-stroke recovery.

**Novelty and Significance:** *What Is Known?:*

- Infiltrated monocyte-derived macrophages (Mφ) into the post-ischemic brain cause neural inflammation, but they also engage in efferocytosis that promotes tissue repair in the injured CNS.
- Remote ischemic limb conditioning (RIC) changes monocyte composition and enhances functional recovery in experimental brain ischemia.
- The application of RIC is safe, feasible, and tolerable in stroke patients, but clinical outcomes remain inconsistent.

*What New Information Does This Article Contribute?:*

- We provide experimental evidence that RIC modifies peripheral monocyte composition and molecular expression, and leads to favorable changes in debris clearance, structure integrity, transneuronal degeneration, and behavior following stroke.
- Protective effects of RIC disappear in the absence of CD36 in Mφ, suggesting an essential mechanistic role for CD36 in RIC-induced endogenous protective outcomes.
- The current study demonstrates that immune-mediated RIC mechanisms facilitate inflammatory and recovery processes in the injured CNS. Given the challenges in directly manipulating the brain after stroke, the study suggests that RIC is a promising alternative strategy by inducing changes in peripheral monocytes that can influence injury progression and recovery. Moreover, RIC-induced peripheral changes uncovered by this study may serve as biomarkers to establish an optimal RIC protocol.

## Introduction

Stroke is a leading cause of physical disability, but the successful management and treatment of stroke remain to be limited. The difficulties of directly manipulating the brain and accessing agents through BBB are attributed to the limitation. These challenges advocate for the development of alternative strategies, including the use of peripheral systems to influence stroke pathophysiology.

Stroke causes a massive infiltration of peripheral immune cells into the injured brain ^1,2^. Two distinct monocyte subsets, representing anti-inflammatory (Ly6C^Low^/CCR2-) and pro-inflammatory (Ly6C^High^/CCR2+) monocytes, circulate in mouse blood. Ly6C^High^ monocytes enter the brain during stroke, differentiate into disease-associated Mφ (M1φ), and cause neural inflammation. Despite their contribution to tissue damage and infarct development, preventing Ly6C^High^ monocyte entry hinders subsequent stroke recovery ^2–4^. This paradoxical role of Mφ derives from their phenotypic conversion from inflammatory M1φ to reparative “microglia-like” M2φ phenotype in the injury environment ^5–7^. This phenotypic plasticity has contributed to the controversy over the function of Mφ and microglia in the injured brain. Identification of the origin and function of Mφ via *in-situ* tracking of peripheral monocytes in the post-stroke brain showed the sustained contribution of Mφ to stroke pathology and repair from acute and recovery phases of stroke ^6,7^.

In light of the repeated translational failure of neuroprotection-based strategies in stroke, remote ischemic limb conditioning (RIC), a noninvasive paradigm, has emerged as a potential strategy for clinical application in stroke patients ^8–11^. The conditioning-induced protective mechanisms shared among pre-, per-, and post-ischemic applications have been associated with proinflammatory mediators such as iNOS, cytokines, chemokine, toll-like receptors (TLRs), and high mobility group box 1 protein (HMGB1) ^12–17^. Several mechanisms, including increased blood flow, changes in neural and circulatory factors, and immune cells, specifically monocytes have been suggested to mediate cross organ protection away from the site of ischemic conditioning (e.g., limb) ^2,18–22^. The RIC-induced cross-organ protection from preclinical studies promoted clinical investigation in stroke. While RIC is safe, feasible, and tolerable in patients with acute ischemic stroke treated with thrombectomy in patients ^23^, the effectiveness of the RIC for acute stroke remains uncertain due to the variability of doses and intervals of RIC application ^24–26^. Thus, optimization of RIC protocols requires an improved mechanistic understanding of how RIC enhances endogenous protection. Previous studies showed that RIC-induced peripheral monocyte shift to a pro-inflammatory status promoted stroke recovery in mice, and that the absence of the shift in monocyte phenotypes can abolish the RIC-enhanced functional benefit ^2^. Still to be delineated are the molecular and cellular events that temporally connect RIC-induced alteration in monocyte composition to behavior outcomes and how acute changes in the blood foster long-term functional improvement.

Mapping longitudinal transcriptome changes in the post-ischemic brain showed that the most perturbed and persistently upregulated genes in the ipsilateral hemisphere were related to neuroinflammation and the clearance of cellular debris (efferocytosis) ^27^. The promotion of efferocytosis is mediated by an orchestrated interaction between danger signals and phagocytic receptors that drive the conversion of disease-associated Mφ to reparative “microglia like” Mφ in CNS environments ^5,28–32^. Mechanistically, efferocytosis-induced changes in metabolism, resulting in greater lactate production, promoted proliferation of reparative Mφ, the key process for tissue resolution ^33,34^. In stroke, efferocytosis contributes to hematoma and injury resolution in the post-ischemic brain ^35–37^. CD36, an innate pattern recognition receptor, is expressed in various cell types, including monocytes and Mφ (Mo/Mφ), microvascular endothelial cells, microglia, platelets, and epithelial cells in many tissues ^38,39^. CD36 elicits inflammatory innate host responses and clears apoptotic cells and debris for tissue resolution ^40–44^. Based on this functional duality, we hypothesize that CD36 mediates RIC-associated molecular and cellular changes that promote functional recovery in chronic stroke. Here, we show that Mo/Mφ-CD36 mediates RIC-induced monocyte shift to a pro-inflammatory state and efferocytosis, and that this shift is associated with an attenuation of transneuronal degeneration and improvement in stroke recovery. These findings suggest that the upregulation of peripheral CD36 is a potential approach to mimic RIC effects.

## Methods

### Data Availability

The data of the article can be obtained by sending a reasonable request to the corresponding author.

**Detailed methods and the Major Resources Table are provided in the Supplemental material.**

## Results

### RIC increases monocyte entry and enhances Mφ efferocytosis in the post-ischemic brain

We previously showed that RIC applied 2h after stroke (post-conditioning) induces a shift in monocytes towards a pro-inflammatory (Ly6C^High^/CCR2+) subset during the acute phase of stroke, and that this shift is necessary for enhanced stroke recovery ^2^. This previous study demonstrated RIC-induced monocyte composition changes in the blood were important for enhanced functional recovery, but the important question of how a monocyte shift in the acute phase resulted in long-term behavior benefit remained unanswered. Since clearing apoptotic cells and cellular debris by Mφ is crucial for injury resolution and tissue repair, the present study asked whether RIC-enhanced efferocytosis by monocyte-derived Mφ in the injured brain is a key biological response that connects the temporally distant events of acute cellular change and improved behavior outcome.

To address whether RIC enhanced efferocytosis, mice were subjected to 30 min transient MCAO and received either sham conditioning (Sham) or RIC at 2h after stroke. At 3d post-ischemia, brain immune cells were harvested from each hemisphere and analyzed by flow cytometry to assess efferocytosis activity *in vivo* (Figure 1A). Mφ and microglia were identified as a [CD11b+/CD45+/Lin-] population by excluding [Lin+] T- and B-cells, NK cells, and neutrophils (Figure 1B). Only a [CD45^Low^] subset was present in the contralateral hemisphere, whereas the ipsilateral hemisphere contained both [CD45^High^] and [CD4 ^Low^] subsets (Figure 1B)^45^. RIC did not increase the number of CD45+ cells in the contralateral hemisphere, but it significantly increased the CD45+ population in the ipsilateral hemisphere. Together, these findings show RIC promotes the infiltration of circulating monocytes into the injured hemisphere and these cells adopt a [CD45^High^] Mφ phenotype (Figure 1C).

**Figure 1.**
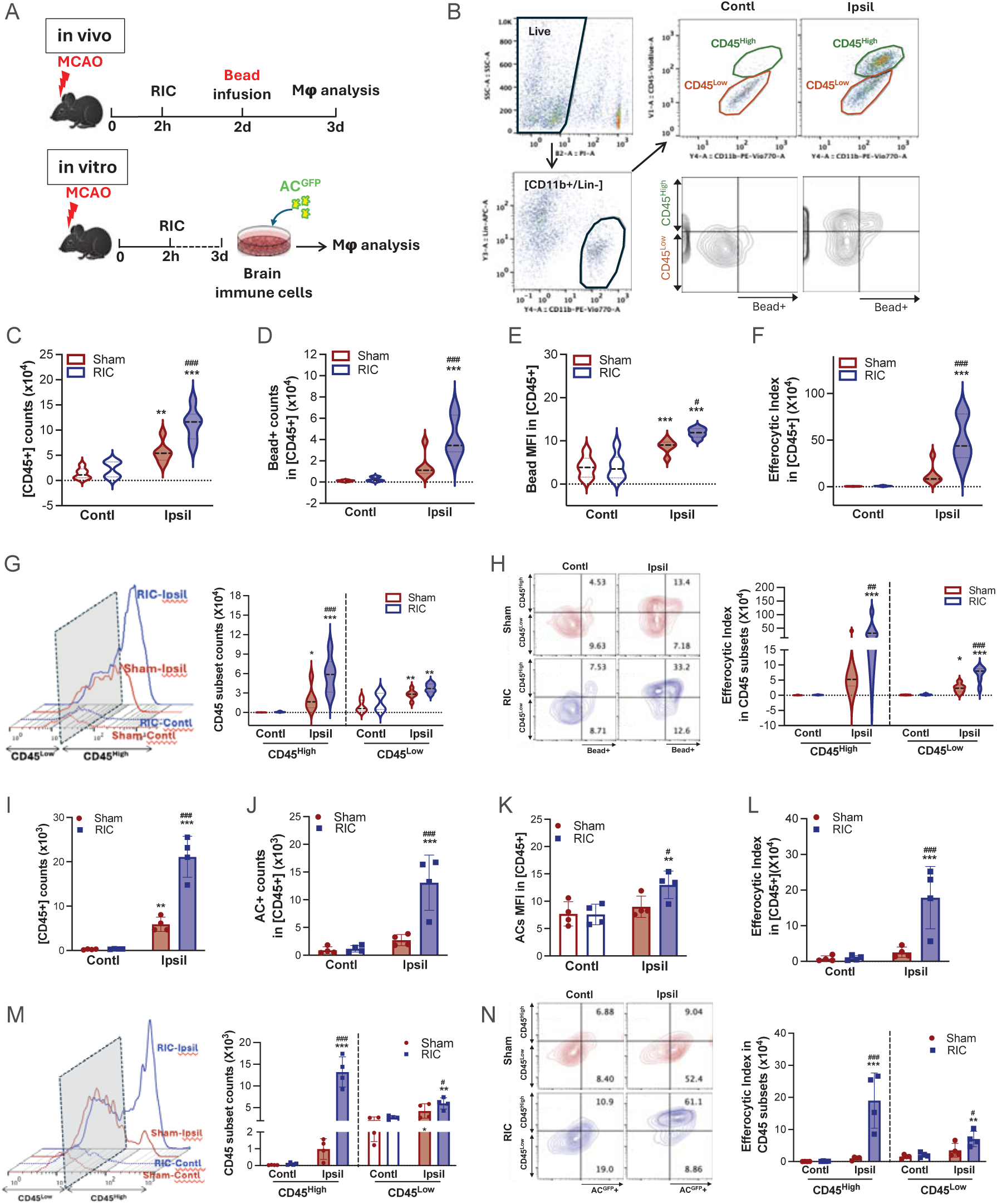
RIC increases monocyte entry and enhances. **M**φ **efferocytosis in the post-ischemic brain. A,** Experimental timeline. Mice were subjected to 30 min MCAO and RIC was applied 2 h after reperfusion. Immune cells isolated from each hemisphere were analyzed at 3d post-MCAO. For *in vivo* efferocytosis assay, beads^580/605^ were infused retro-orbitally into the mice at 2d post-MCAO. For *in vitro* efferocytosis assessment, isolated brain immune cells were cultured for 2h (2x10^5^/well). GFP-tagged apoptotic cells (AC^GFP+^) were then added (brain immune cells: AC^GFP+^=1:5) and co-cultured for 1h. Analysis measured the number of AC^GFP^+ cells within the [CD45+/CD11b+/Lin-] immune cells. **B,** Gating of [CD45+/CD11b+/Lin-] Mφ/microglia in the post-ischemic brain to determine bead+ cell number (counts) and mean fluorescent intensity (MFI) of efferocytosis. **C-H,** *In vivo* analyses (n=6/group). Effect of RIC on the number of phagocytes and efferocytosis in [CD45+/CD11b+/Lin-] populations. CD45+ Mφ/microglia count (**C**), Bead+ cell count (**D**), Mean fluorescence intensity (MFI) of bead+ cells (**E**), Efferocytosis index (bead+ counts X bead MFI (**F**), CD45+ Mφ/microglia counts in CD45 subsets (**G**), Efferocytosis index in CD45 subsets (H) **I-N,** *In vitro* analyses (n=4/group) of RIC effects on the number of phagocytes and efferocytosis in [CD45+/CD11b+/Lin-] populations. CD45+ Mφs/microglia count (**I**), AC+ cell count (**J**), Mean fluorescence intensity (MFI) of AC+ cells (**K**), Efferocytosis index (AC+ counts X AC MFI (**L**), CD45+ Mφ/microglia counts in CD45 subsets (**M**), Efferocytosis index in CD45 subsets (**N**). Statistical significance was assessed with two-way ANOVA followed by *post hoc* Fisher’s LSD test. ^*, **,***^ *p* < 0.05, 0.01, 0.001 Contl vs. Ipsil (Effect of stroke); ^#, ##, ###^ *p* < 0.05, 0.01, 0.001 Sham vs. RIC (Effect of RIC). Abbreviations: Contl (contralateral); Ipsil (ipsilateral); Sham (sham conditioning); RIC (remote ischemic conditioning).

To assess *in vivo* Mφ efferocytotic activity in the brain, fluorescence beads (beads^580/605^) were infused intravenously at 2d after MCAO (Figure 1A). The beads readily cross the blood-brain barrier and can be phagocytosed by brain immune cells. At 3d post-stroke, brain cells were dissociated and analyzed by cytometry to count the [CD45+/CD11b+/Lin-] cells containing beads (Figure 1B). Our previous study showed that Mφs, not microglia, are the major phagocytes in the post-ischemic brain and that circulating monocytes do not phagocytose beads ^6^, which indicates that bead+ CD45+ cells in the post-ischemic brain in the current study are predominantly Mφs engaging efferocytosis. Flow cytometry analysis of the CD45+ population in the current study showed that RIC significantly increased the number of beads+ CD45+ cells and bead intensity (mean fluorescent intensity, MFI) in the ipsilateral hemisphere (Figure 1D-E). Accordingly, the efferocytosis index (counts x MFI) was significantly and selectively increased in the RIC group (Figure 1F).

Additional flow cytometry analyses addressed whether there were differences in trafficking and efferocytosis levels between the CD45^High^ and CD45^Low^ subsets. RIC significantly increased the number of CD45^High^ cells in ipsilateral hemispheres at 3d post-stroke (Figure 1G). RIC increased efferocytosis levels in both CD45 subsets at 3d post-stroke (Figure S2A, B), but the magnitude of efferocytosis activity was greater in the CD45^High^ subset (Figure 1H). Given that CD45^High^ Mφs undergo phenotype changes to a “microglia-like” CD45^Low^ phenotype in the post-ischemic brain ^6,7^ and injury environment ^5,28^, the increase in both the number and efferocytosis levels of CD45^Low^ cells in the current study were likely CD45^High^ Mφ that have adopted the CD45^Low^ reparative phenotype.

To test whether the RIC could also enhance efferocytosis levels in an *in vitro* model system, we measured the efferocytosis activity of brain immune cells isolated 3d post-MCAO that were co-cultured with apoptotic cells (ACs; Figure 1A). To generate ACs, isolated cultured splenocytes were placed under UV light for 30 min ^46^. Splenocytes cultures were also labelled with GFP using GFP-Cell Linker (>99% GFP labelling efficiency and ∼70% were ACs; Figure S3). After 1h of co-culture with ACs, brain immune cells [CD45+/CD11b+/Lin-] isolated from the ipsilateral hemisphere of stroked mice treated with RIC contained a significantly higher number of cells that contained significantly higher of AC^GFP+^ cells when compared to either to brain immune cells isolated from the contralateral hemisphere or from sham-treated mice. This confirmed the *in vivo* findings that RIC enhances immune cells within ipsilateral hemisphere from mice to have significantly elevated levels of efferocytosis and MFI (Figure 1J-L).

Additional analyses of the cultured brain immune cells also showed cells isolated from stroked mice treated with RIC contained a significantly higher number of CD45^High^ and CD45^Low^ cells in the ipsilateral hemisphere (Figure 1M). Moreover, RIC significantly enhanced efferocytosis activities in both CD45 subsets (Figure 1N), and the enhancement is mainly from the increased number of AC^GFP+^ Mφs and not from an increased efferocytosis capacity (MFI) (Figure S4A, B). Taken together, the *in vivo* and *in vitro* findings show that RIC enhances monocyte trafficking into the injured hemisphere and increases the Mφ population and efferocytosis activity in the post-ischemic brain. These effects collectively indicate that RIC can modulate the peripheral immune system to CNS injury and the subsequent repair processes.

### RIC-induced CD36 expression enhances efferocytosis in the post-ischemic brain

Longitudinal mapping of transcriptome changes in the mouse brain for 6 months after stroke showed that the most perturbed and persistently upregulated genes are related to neuroinflammation and pattern recognition of clearance of cellular debris and apoptotic cells (NCBI SRS [National Center for Biotechnology Information Short Read Archive], accession number: PRJNA525413) ^27^ (Figure S5A,B). CD36 is a class B scavenger receptor involved in innate host immunity and efferocytosis ^40–44,47^ that has elevated expression for months after stroke (Figure S5B), suggesting that CD36 may be an effector molecule for RIC-induced immune modulation in Mφs.

To establish whether CD36 mediated, at least in part, the effects RIC on the peripheral immune system following stroke and/or the subsequent repair processes, we initially addressed whether RIC impacted CD36 expression levels in [CD45+/CD11b/Lin-] cells in the post-ischemic brain at 3d (Figure 2A). Stroke increased the number of CD36+ cells (Figure 2B) and CD36 expression levels (MFI, Figure 2C) for both sham and RIC groups in the ipsilateral hemisphere. Compared to the sham group, the increases were significantly larger in the RIC group (indicated by #, Figure 2B, C). The RIC-induced increase in CD36+ cells occurred in both CD45 subsets, although the increase in the CD45^High^ subset was significantly higher (3.1 vs 1.5-fold increase for CD45^High^ and CD45^Low^, respectively; Figure 2E). RIC increased CD36 expression (MFI) only in the CD45^High^ subset (Figure 2F).

**Figure 2.**
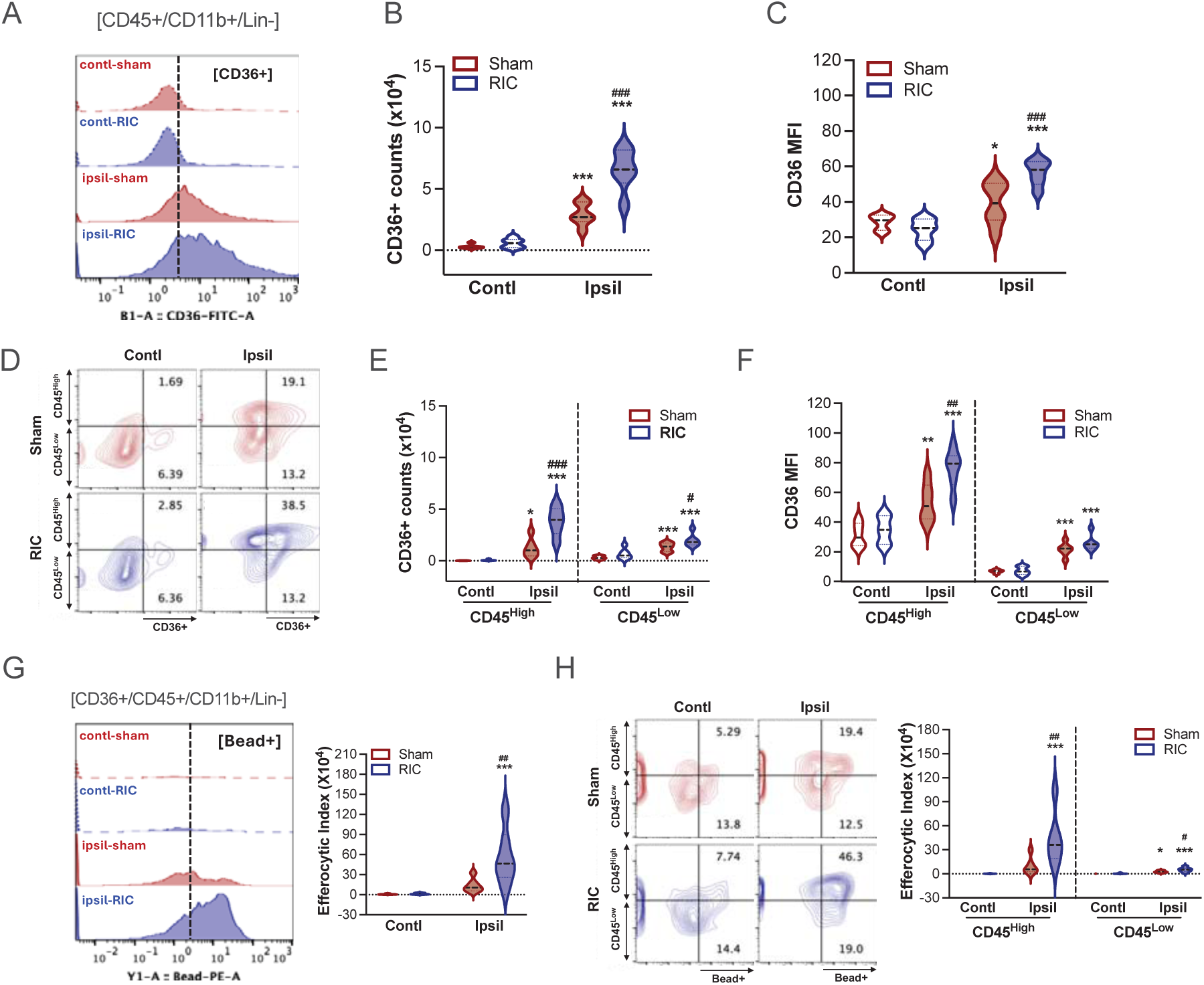
RIC-induced CD36 expression enhances efferocytosis in the post-ischemic brain. Brain immune cells were isolated at 3d post-stroke from the contralateral and ipsilateral hemispheres of sham or RIC animals. **A-C,** Analyses of CD36 expression in [CD45+/CD11b+/Lin-] cells. CD36 flow histogram in sham and RIC animals (**A**), number of CD36+ cells (**B**), and CD36 expression (by mean fluorescent intensity, MFI) in total CD45+ cells. **D-F,** Analyses of CD36 expression in CD45^High^ and CD45^Low^ subsets in [CD45+/CD11b+/Lin-] populations. Represent flow diagram (D), number of CD36+ cells (E), and CD36 expression (F) in CD45 subsets. **G-H,** Efferocytosis assay in CD36+ cells in [CD45+/CD11b+/Lin-] cells (**G**) and CD45 subsets (**H**). n=6/group. Statistical significance was assessed with two-way ANOVA followed by *post hoc* Fisher’s LSD test. ^*, **, ***^ *p* < 0.05, 0.01, 0.001 Contl vs. Ipsil (Effect of stroke); ^#, ##, ###^ *p* < 0.05, 0.01, 0.001 Sham vs. RIC (Effect of RIC). Abbreviations: Contl (contralateral); Ipsil (ipsilateral); Sham (sham conditioning); RIC (remote ischemic conditioning).

Because CD36 expression in Mφ positively correlates with phagocytosis activity ^37^, we tested whether RIC modulated CD36-mediated efferocytosis. Mice were infused with beads^580/605^ at 2d post-stroke, and then the [CD45+/CD11b+/Lin-] cells in the brain were isolated and analyzed at 3d post-ischemia (Figure 1A). For CD36-/CD45+ cells, RIC did not impact either the number of bead+ cells, the bead MFI, or efferocytosis activity when compared between either contralateral vs. ipsilateral hemispheres (Figure S6 A-C) or between CD45^High^ and CD45^Low^ subsets (Figure S6 D-F). By contrast, RIC significantly increased bead+ and bead MFI (Figure S7A, B), as well as efferocytosis (Figure 2G), for CD36+ cells within the ipsilateral hemisphere, when compared to the contralateral hemisphere (indicated by *) and compared to the sham (indicated by #). RIC significantly increased bead+ cells and bead MFI in both CD45^High^ and CD45^Low^ subsets in the ipsilateral hemisphere (Figure S7C, D), which resulted in significantly higher efferocytosis levels, especially for the CD45^High^ subset (Figure 2H). Together, these results show that RIC augments stroke-induced CD36 expression and CD36-mediated efferocytosis in the injured brain. Moreover, the higher levels of CD36+/CD45^High^ cells in the ipsilateral hemisphere of RIC-treated mice after stroke suggests that peripherally derived CD36+ Mφ have a major role in the RIC-enhanced efferocytosis in the post-ischemic brain.

### CD36 mediates a RIC-induced monocyte shift to a pro-inflammatory state

Since previous work showed that the periphery is the major source of CD36-expressing cells in the post-ischemic brain^48^, we next addressed whether RIC triggers an upregulation of CD36 expression in circulating monocytes before they enter the injured brain. To determine the effect of RIC on CD36 expression in monocytes, RIC was applied to naïve mice without stroke to avoid stroke-induced monocyte mobilization to the injured brain. The blood from naïve WT mice with either RIC or sham was collected at 1d post-RIC, and CD36 expression in monocytes was analyzed by flow cytometry (Figure 3A, bottom left). RIC did not alter number of either total or CD36+ monocytes (Figure 3B, C), but it did significantly increase CD36 expression levels in monocytes (Figure 3D). Circulating monocytes in mice are a heterogeneous population of pro-inflammatory Ly6C^High^ [CCR2+] and anti-inflammatory Ly6C^Low^ [CCR2-] subsets. Given that RIC shifts monocyte compositions to a proinflammatory state (increased Ly6C^High/Low^ ratio), and this shift is necessary to promote stroke recovery ^2^, the analysis of peripheral monocytes in naïve mice in this current study showed that RIC increased Ly6C^High^ monocytes without altering an Ly6C^Low^ subset (Figure 3E, left), and generated an overall shift towards a proinflammatory state (Figure 3E, right). Moreover, RIC-enhanced CD36 expression occurred selectively in Ly6C^High^ monocytes (Figure3 F), which is the subset that enters the post-ischemic brain. The RIC-induced upregulation of CD36 in Ly6C^High^ monocytes thus accounts for the increases in the number of CD36+ monocytes (Fig 2B, E) and the CD36 MFI (Fig 2C, F) in the post-ischemic brain. When compared to CD45^High^ Mφ in the brain after stroke, however, the RIC-mediated increase CD36 levels in circulating Ly6C^High^ monocytes was proportionately less (cf. Figure 2F vs. Figure 3F). These findings indicate RIC triggers an increase in CD36 expression within peripheral monocytes and suggest that the abundance of CD36 ligands in the injured brain ^49–52^ further elevates CD36 expression levels following their infiltration into the ipsilateral hemisphere.

**Figure 3.**
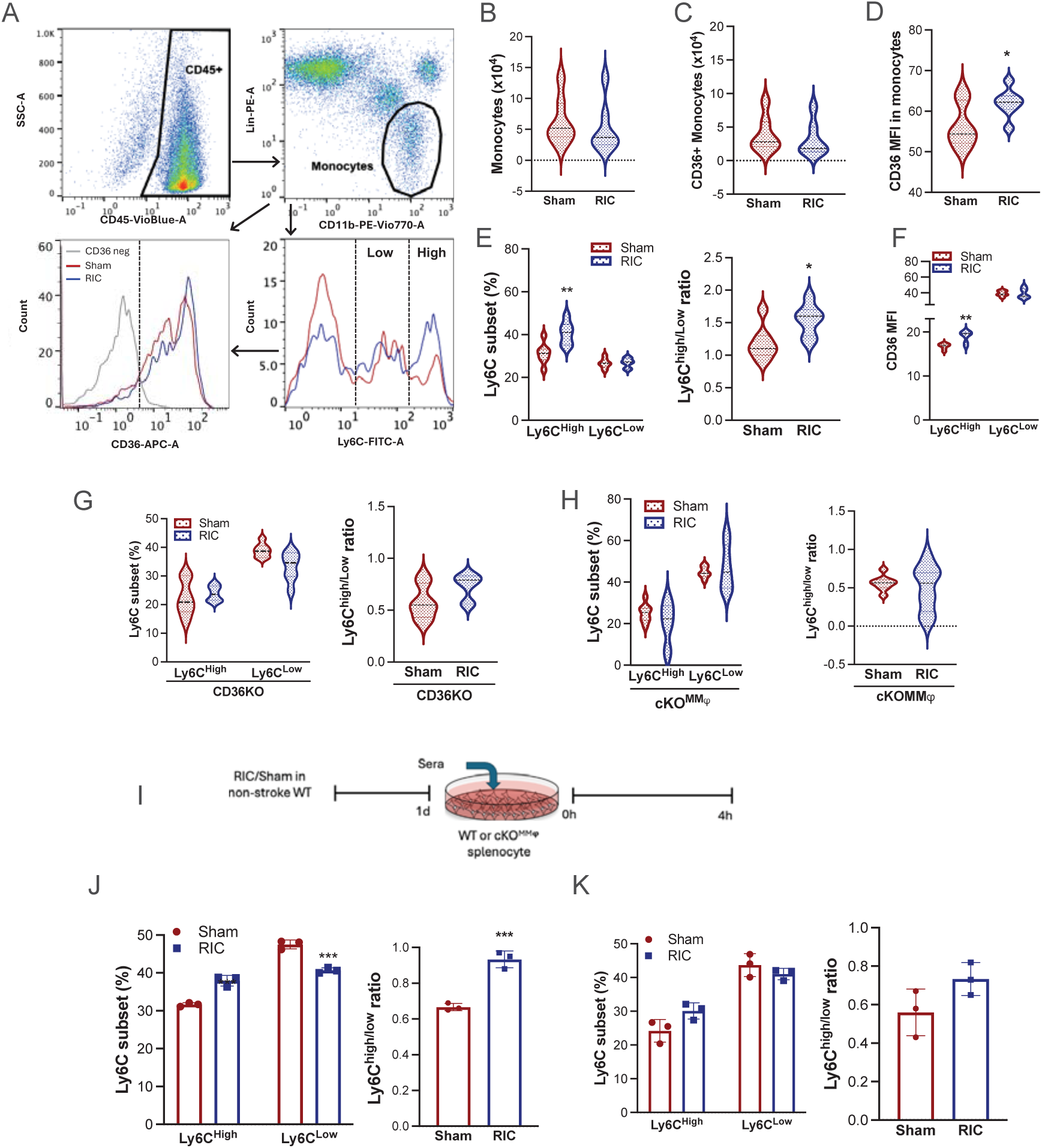
CD36 mediates a RIC-induced monocyte shift to a pro-inflammatory state. The blood of naïve non-stroke mice was analyzed at 1d after sham or RIC. N=7/group **A,** Flow gating strategy in the blood of naïve non-stroke mice 1d after sham or RIC. Histogram shows CD36 expression in monocytes (lower left) and monocyte Ly6C subsets (lower right), N=7/group **B-D,** Quantification of total number of circulating monocytes [CD45+/CD11b+/Lin-], CD36+ monocytes, and CD36 expression (MFI) in monocytes. **E,** Effect of RIC on monocyte subsets, Percentage of Ly6C^High^ and Ly6C^Low^ monocyte subsets (left) and Ly6C^High/Low^ ratio (right) **F,** CD36 MFI in the Ly6C monocyte subsets. **G-H**, Effect of RIC on Ly6C^High^ or Ly6C^Low^ monocyte subsets (left) and Ly6C^High/Low^ ratio (right) in CD36KO mice (G, n=7) and cKO^MMφ^ mice (H, n=8). **I**, Sera were harvested 1d after sham or RIC in naïve WT mice. Splenocytes were incubated with Sham or RIC-sera for 4 hours and monocyte subsets were analyzed. **J**-**K**, Percentage of Ly6C^High^ or Ly6C^Low^ monocyte subsets (left) and the Ly6C ^High/Low^ ratio (right) in WT splenocytes (**J**) and in cKO^MMφ^ splenocytes (**K**). T-test: *, **, *** *p*<0.05, 0.01, 0.001 Sham vs. RIC

To establish the necessity of CD36 for the RIC-induced monocyte shift, we analyzed monocytes in mice with a constitutive CD36 deletion (CD36KO). This analysis found that RIC neither induced a change in the number of monocytes nor a shift in subsets (Figure 3G). The interpretation of these findings is confounded, however, by the fact that CD36 expressed in many cell types and tissues. To assess the role of CD36 specifically in monocyte/Mφ, mice with a conditional deletion of CD36 in monocytes/Mφs (cKO^MMφ^) were analyzed (Figure S8). Like the CD36KO mice, an RIC-induced change in either the number of monocytes or shift monocyte subsets was absent in cKO^MMφ^ mice (Figure 3H). The disappearance of the subset shifts in both CD36KO and cKO^MMφ^ mice demonstrates that CD36 expression in monocytes/Mφ is necessary for the RIC-induced shift in monocyte inflammatory states.

The requirement for CD36 expression in monocytes/Mφ was also tested *in vitro* by incubating splenocytes with sera from RIC or sham mice (Figure 3I). When compared to cultures treated with sera from sham mice, WT splenocytes treated with sera from RIC-treated mice showed a significant increase in the Ly6C^High^ subset and a decrease the Ly6C^Low^ subset, which increased Ly6C^High^/^Low^ ratio (Figure 3J). By contrast, cKO^MMφ^ splenocytes treated with sera from RIC-treated mice did not induce shifts in monocyte subsets (Figure 3K). Together with the *in vivo* findings, these findings show that CD36 is required for RIC-induced changes in both monocyte composition and the shift in monocyte inflammatory state.

### cKO^MMφ^ mice delay stroke recovery

The shift monocyte in inflammatory state is required for promoting recovery in chronic stroke ^2^ and this shift is abrogated in cKO^MMφ^ mice. We next examined whether the loss of CD36 in Mφ impacted stroke outcomes. Compared to the WT genotype, cKO^MMφ^ mice had similar mortality during an acute phase of stroke, comparable stroke-induced body weight loss, and similar infarct size when assessed at 3d post-ischemia (Figure 4A-C). The number of cells in the CD45^High^ subset within the ipsilateral hemisphere was also similar, suggesting that monocyte trafficking between the WT and cKO^MMφ^ genotypes was comparable (Figure 4D). The number of the CD45^Low^ cells, however, was significantly larger in both the ipsilateral and contralateral hemispheres of cKO^MMφ^ mice (Figure 4D), suggesting that the loss of CD36 in Mφ/monocytes influences the microglia populations and/or Mφ that adopt a “microglia-like” CD45^Low^ phenotype.

**Figure 4.**
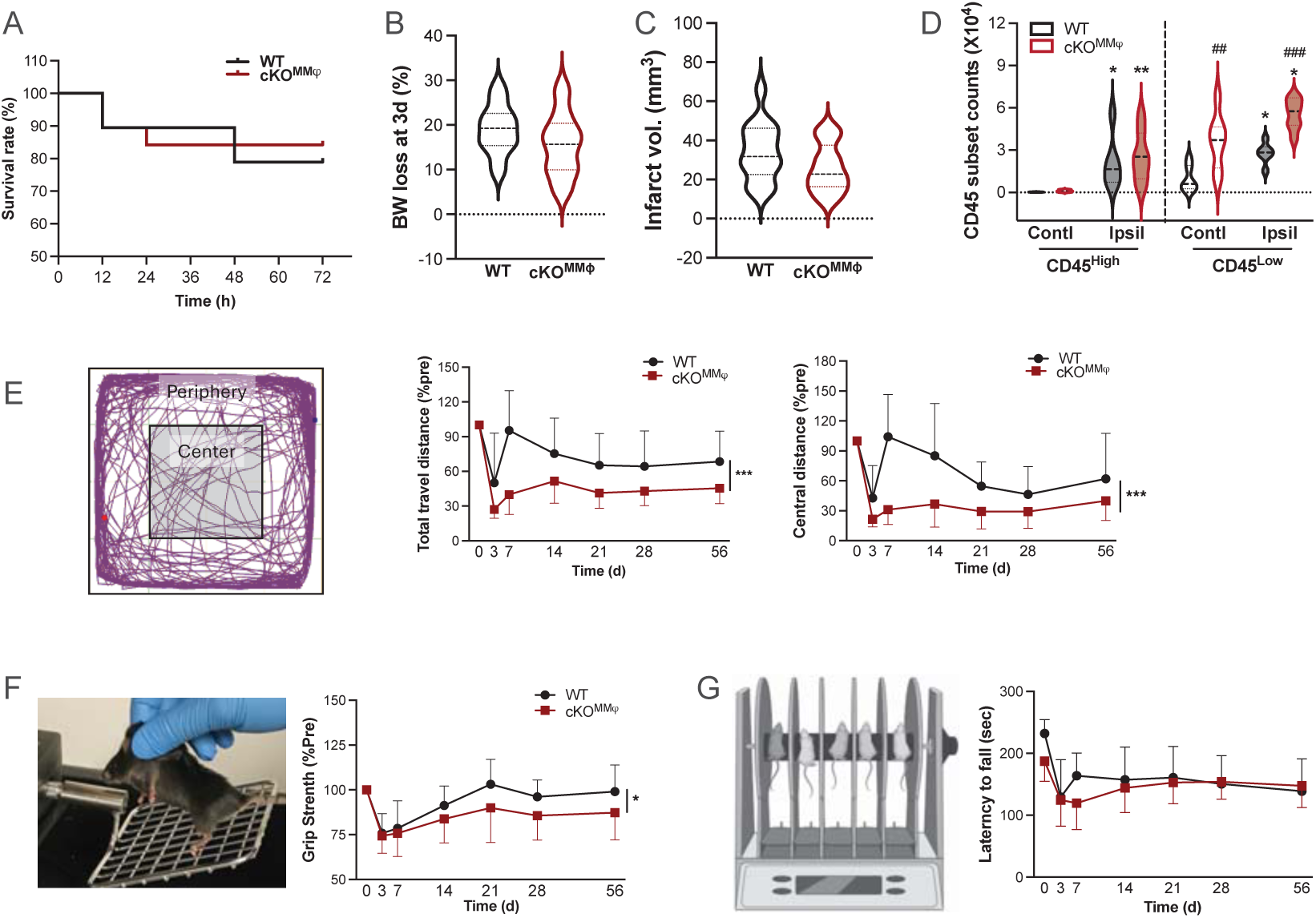
cKO^MMφ mice delay stroke recovery.^ **A**, Assessment of survival rate in WT (CD36^fl/fl^) and cKO^MMφ^ mice during an acute phase of stroke, n=19-20/group at a starting point. **B**, Assessment of body weight loss (% prestrike baseline) at 3d post-MCAO, n=16 **C**, Indirect (edema corrected) infarct volume at 3d post-MCAO, n=14-16/group. **D**, Flow analyses of CD45 subset in [CD11b+/Lin-] populations, n=6-7/group. Abbreviations: Contl (contralateral); Ipsil (ipsilateral). **E-G**, Longitudinal post-stroke behavior test in WT or cKO^MMφ^ mice (n=7-9/group) Open field test to assess total travels distance and distance in the center zone against pre-baseline (% pre, **E**), Grip strength against pre-baseline (% pre, **F**), and Rotarod (**G**). Two-way ANOVA: *, *** *p* < 0.05, 0.001 WT vs. cKO^MMφ^ (effect of genotype)

Despite similar acute outcomes, cKO^MMφ^ mice displayed greater motor deficits than the WT mice. Longitudinal open field tests showed that cKO^MMφ^ mice underperformed in traveling distance and less time in the center in an open field when compared to their pre-stroke baselines (Figure 4E). The cKO^MMφ^ mice also reduced grip strength of the hind limbs when compared to WT mice (Figure 4F). By contrast, we did not observe a difference in rotarod performance between genotypes (Figure 4G). Unlike open field and grip strength tests, which showed similar pre-stroke baseline between genotypes, rotarod showed a significant difference in baseline between the genotypes (232.1 ± 29.4sec vs 187.4± 45.4sec for WT and cKO^MMφ^, respectively; n=9-10/genotype, *p*<0.05). This suggests that cKO^MMφ^ mice had impaired motor learning ability during pretraining, and this may mask differences in recovery from stroke-induced motor deficits. The overall delayed stroke recovery in cKO^MMφ^ mice reveals the importance of CD36-expressing Mφ for functional recovery in stroke.

### Stroke-induced efferocytosis is reduced in cKO^MMφ^ mice

To address that CD36 was necessary for RIC-enhanced efferocytosis, we determined the effect of Mφ-CD36 deletion on efferocytosis in the post-ischemic brain. WT and cKO^MMφ^ brain immune cells isolated at 3d post-ischemia were incubated AC^GFP+^ (Figure 1A, *in vitro*). Since the monocyte infiltration, indicated by the CD45^High^ subset, was comparable between WT and cKO^MMφ^ mice (Figure 4D), the same number of brain immune cells (2x10^5^/well) from these mice were cultured for efferocytosis assay. For WT cells, stroke increased the number of AC^GFP+^ cells and their MFI (Figure 5B, C), leading to a significantly enhanced efferocytosis activities (Figure 5D). For cKO^MMφ^ cells, despite stroke-induced increases in the number of AC^GFP+^ cells and MFI (Figure 5B, C), the efferocytosis index in cKO^MMφ^ did not reach significance, and significantly lower compared to WT cells (Figure 5D, indicated by #). These findings show that stroke selectively enhances efferocytosis in the ipsilateral hemisphere of WT, but not in cKO^MMφ^ mice.

**Figure 5.**
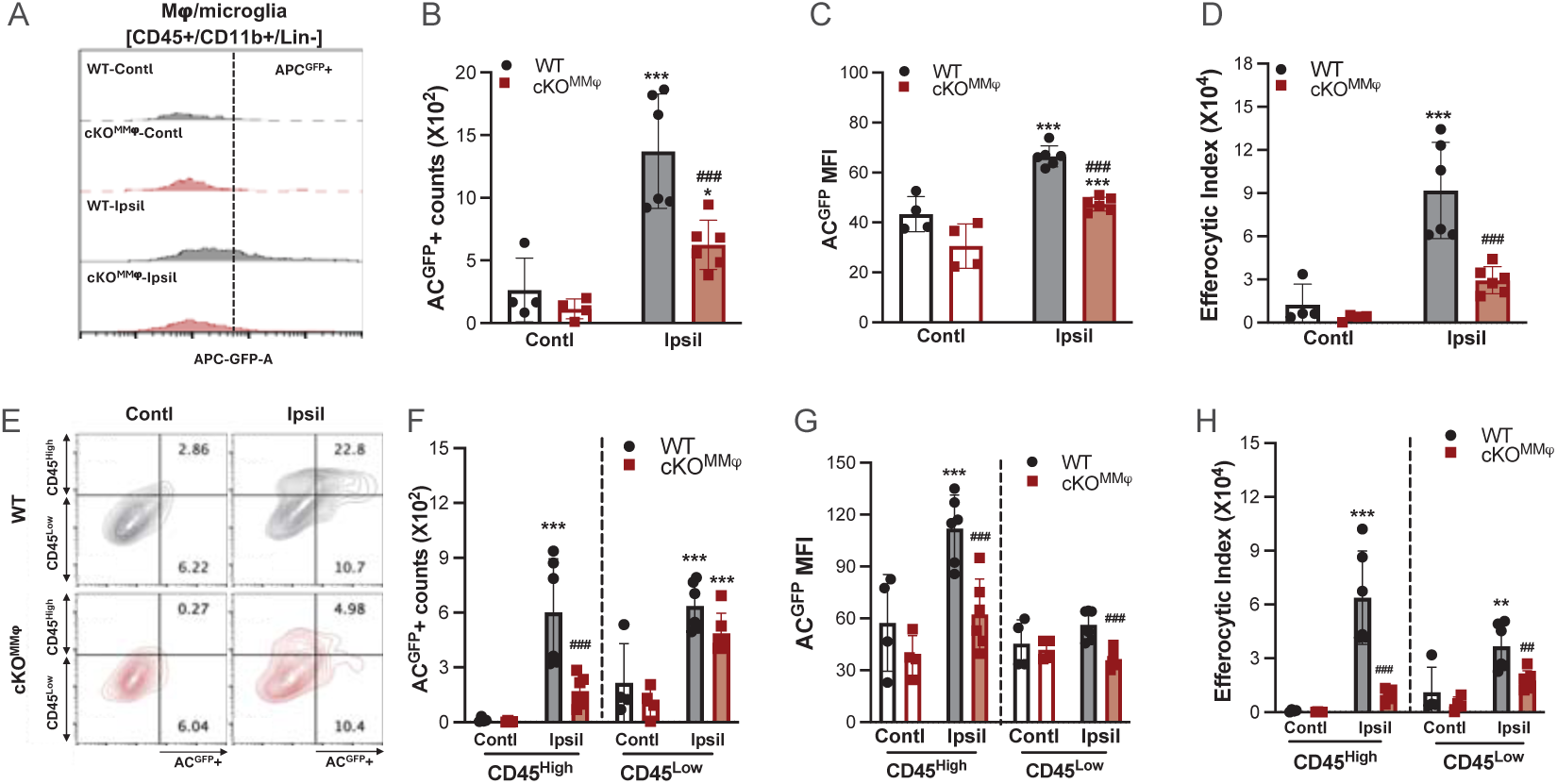
Stroke-induced efferocytosis is reduced in the cKO^MMφ^ mice. GFP-tagged apoptotic cells (AC^GFP+^) were added to the cultures (brain cells: ACs, 1:5) for 1h and analyzed AC^GFP^+ cells in [CD45+/CD11b+/Lin-] Mφs/microglia. **A,** Flow histogram of AC^GFP^+ cells in Mφs/microglia. **B-D**, Quantification of AC^GFP+^-Mφs/microglia count (**B**), AC^GFP+^-Mφs/microglia mean fluorescence intensity (**C**), Efferocytosis index (AC^GFP+^counts x AC^GFP+^MFI (**D**), **G** Flow contour plot of AC^GFP^+ CD45^High^ and CD45^Low^ Mφs/microglia. **F-H,** Quantification of AC^GFP+^ counts in CD45 subsets (**F**), AC^GFP+^ MFI in CD45 subsets (**G**), Efferocytosis index (AC^GFP+^ counts x AC^GFP+^MFI) in CD45 subsets (**H**). Abbreviations: Contl (Contralateral); Ipsil (Ipsilateral). Two-way ANOVA followed by post-hoc Fisher’s LSD test. ^**, ***^ *p* < 0.01, 0.001 Contl vs. Ipsil (Effect of stroke); ^##, ###^ *p* < 0.01, 0.001 WT vs. cKO^MMφ^ (Effect of genotype).

Additional analyses of the CD45 subsets within cKO^MMφ^ brain immune cells showed reduced efferocytosis in both subsets, but a greater relative reduction in CD45^High^ subset (Figure 5H). The reduction in the CD45^High^ subset derived from a reduction in the number of AC^GFP+^ cells and MFI (Figure 5F, G), whereas the reduction in the CD45^Low^ subset resulted from reduced MFI (Figure 5G). Taken together, these findings show CD36 expression in Mφ is required for stroke-induced efferocytosis in the ipsilateral hemisphere.

### CD36 mediates RIC-enhanced efferocytosis in the post-ischemic brain

The required monocyte shift for RIC-promoted stroke recovery ^2^ as well as the contribution of CD36 expression in Mφ for the monocyte phenotype shift and efferocytosis, together suggest that Mφ-CD36 mediates RIC-enhanced efferocytosis in the post-ischemic brain. Thus, we used cKO^MMφ^ mice to address whether CD36 was required for RIC-enhanced efferocytosis. Unlike RIC-enhanced efferocytosis in WT (C57) mice (Figure 1C-F), applying RIC to cKO^MMφ^ mice did not increase either the number of Mφ/microglia (Figure 6A) or efferocytosis activity (bead+ count, MFI, and efferocytosis index; Figure 6B-D). Analysis of CD45 subset showed that RIC significantly increased efferocytosis in CD45^High^ (and CD45^Low^) subsets in WT mice (Figure 1H), but RIC with cKO^MMφ^ mice had no effect on the number of CD45^High^ and CD45^Low^ cells (Figure 6E), efferocytotic cells, MFI (Figure S9 A, B) or efferocytosis activities (Figure 6F) in both CD45 subsets. Similar findings were seen with cultured brain immune cells from cKO^MMφ^ mice, where RIC had no effect on the number of Mφ/microglia (Figure 6G) and efferocytosis activities (Figure 6H-J). Moreover, RIC did not influence the number of Mφ/microglia (Figure 6K) and efferocytosis activities in CD45 subsets (Figure S9 C, D & Figure 6L). Given that the CD45^High^ subset represents infiltrating monocyte-derived Mφ into the injured brain, the absence of the RIC effects in cKO^MMφ^ mice reveals the importance of CD36 for the effect of RIC and the critical role that Mφ-CD36 have in mediating RIC-enhanced efferocytosis in the injured CNS.

**Figure 6.**
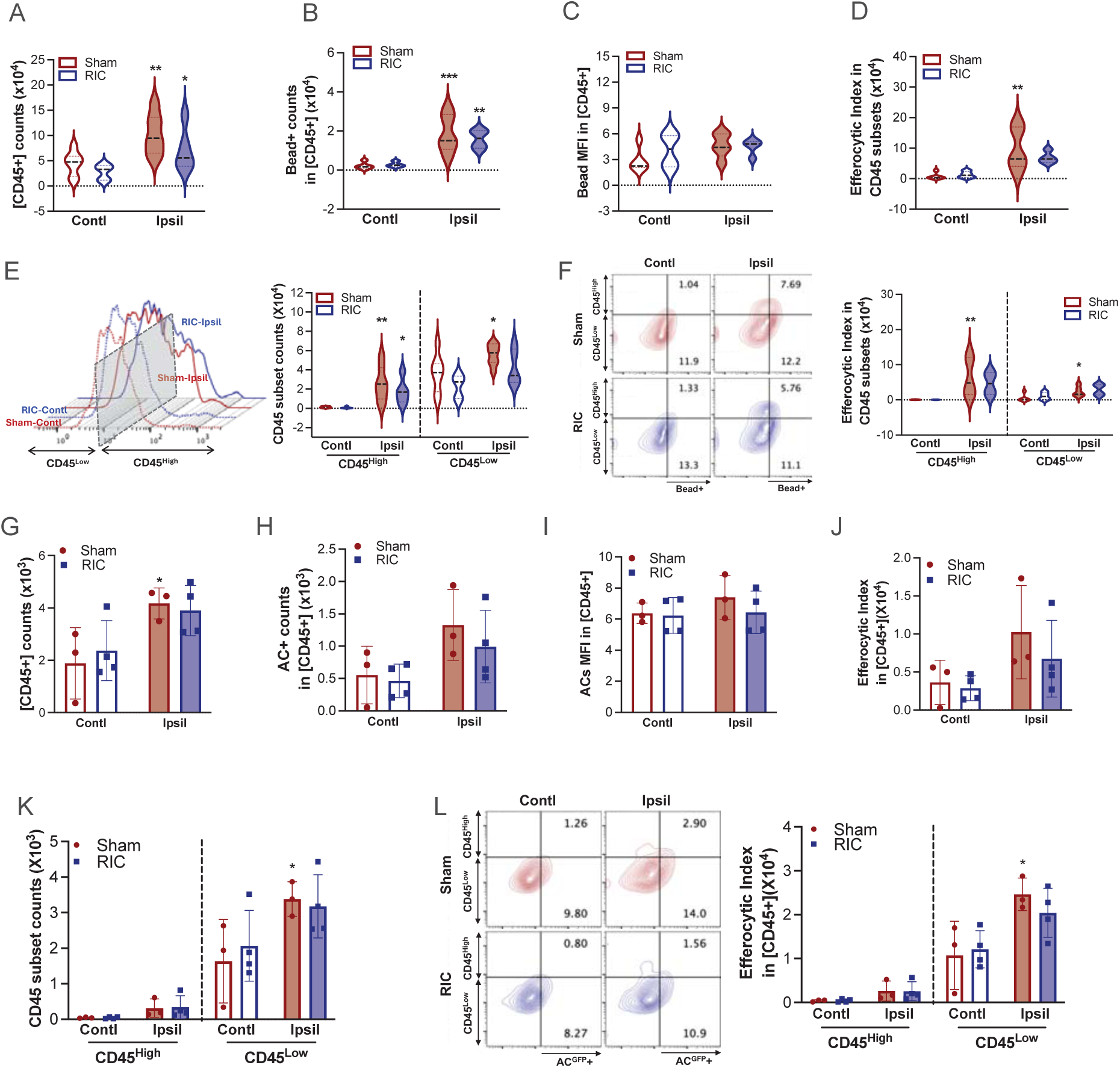
CD36 mediates RIC-enhanced efferocytosis in the post-ischemic brain. cKO^MMφ^ mice with 30 min MCAO were subjected to Sham or RIC at 2h post-MCAO. Beads^580/605^ were infused retro-orbitally at 2d post-MCAO and immune cells isolated from each hemisphere were analyzed at 3d. **A**, Total [CD45+/CD11b+/Lin-] Mφ/microglia counts in cKO^MMφ^ mice, n=5-7/group. **B-D**, *In vivo* efferocytosis assay, n=4-5/group. The number of Beads^580/605^+ cells in cKO^MMφ^ Mφ/microglia (B), Beads^580/605^+ mean fluorescence intensity (MFI) (C), Efferocytosis index (Beads^580/605^+ counts x Beads^580/605^+ MFI) (D). **E**, Representative histogram for total [CD45+/CD11b+/Lin-] Mφ/microglia number in CD45 subsets, n=5-7/group. **F**, Flow contour plot of cKO^MMφ^ mice for the number of Beads^580/605^+ cells in CD45^High^ and CD45^Low^ Mφ/microglia. n=4-5/group. **G-L,** *In vitro* analyses (n=4/group) of RIC effects on the number of phagocytes and efferocytosis assay in cKO^MMφ^ mice. **G**, [CD45+/CD11b+/Lin-] Mφs/microglia count. **H,** AC+ Mφs/microglia count. **I**, Mean fluorescence intensity (MFI) of AC+ cells. **J,** Efferocytosis index (AC+ counts x AC+ MFI, **K**, CD45^High^ and CD45^Low^ Mφs/microglia count, **L**, Flow contour plot of cKO^MMφ^ mice for AC+ cells and efferocytosis index in CD45^High^ and CD45^Low^ Mφs/microglia. Contl, Contralateral; Ipsil, Ipsilateral; Sham, Sham conditioning; RIC, Remote ischemic conditioning Statistical significance was assessed with two-way ANOVA followed by *post hoc* Fisher’s LSD test. ^*, **, ***^ *p* < 0.05, 0.01, 0.001 Contl vs. Ipsil (Effect of stroke)

### RIC attenuates dopaminergic neuron loss in substantia nigra in chronic stroke

MCAO-induced striatal and cortical infarction causes progressive atrophy in the primary infarcted tissue and remote areas ^53–57^. The progressive brain atrophy is accompanied by sustained immune activation and continuous infiltration of peripheral immune cells into primary and delayed entry to secondary injury sites in the brain ^7,27^. Therefore, we investigated the effect of RIC on the progression of brain atrophy by analyzing brain volume in sub-regions at 2 months post-ischemia. The ischemic stroke in sham-conditioned animals showed brain atrophy throughout the ipsilateral hemisphere and primary injury sites, including the striatum and cortex (Figure S10 A-C), as well as the thalamus, which receives input from the striatum and projects to the cortex (Figure S10D). Other remote areas, however, such as the hypothalamus and hippocampus, were not impacted (Figure. S10 E, F) ^58^. Despite the previous study showing RIC enhances stroke recovery ^2^, the subregion volume assessment in the current study showed that RIC did not attenuate stroke-induced brain atrophy in the primary and secondary injury sites. This analysis, however, used tissue at 2 months to calculate brain volume. Inter-animal variability in the glia scar formation surrounding the injured areas could have masked the effect of RIC, and this led us to investigate the impact of RIC in transneuronal degeneration.

The striatum receives dopaminergic input from tyrosine hydroxylase (TH)-expressing neurons in the substantia nigra pars compacta (SNc). Transneuronal degeneration in SNc is only evident during chronic phases of stroke, and not in the acute phase ^7,53,54^. To probe the effect of RIC on the neuronal loss in the SNc, we used transgenic mice that express GFP in TH+ neurons (Tg^TH-GFP+^). 3D whole brain imaging shows GFP expression in the olfactory bulb, SNc, and locus coeruleus, and the horizontal view showed GFP expression of TH+ fibers in the striatum (Figure 7A). Immunostaining with TH antibody confirmed the GFP-expressing cells are TH+ neurons (Supplementary video 1). We used Imaris software to analyze the SNc volume and the number of TH+ cells in the SNc in both hemispheres (Figure 7B). Stroke reduced SNc volume and TH+ neurons in the ipsilateral brain in both sham and RIC mice, but the reduction was significantly attenuated in RIC mice (Figure 7C, D).

**Figure 7.**
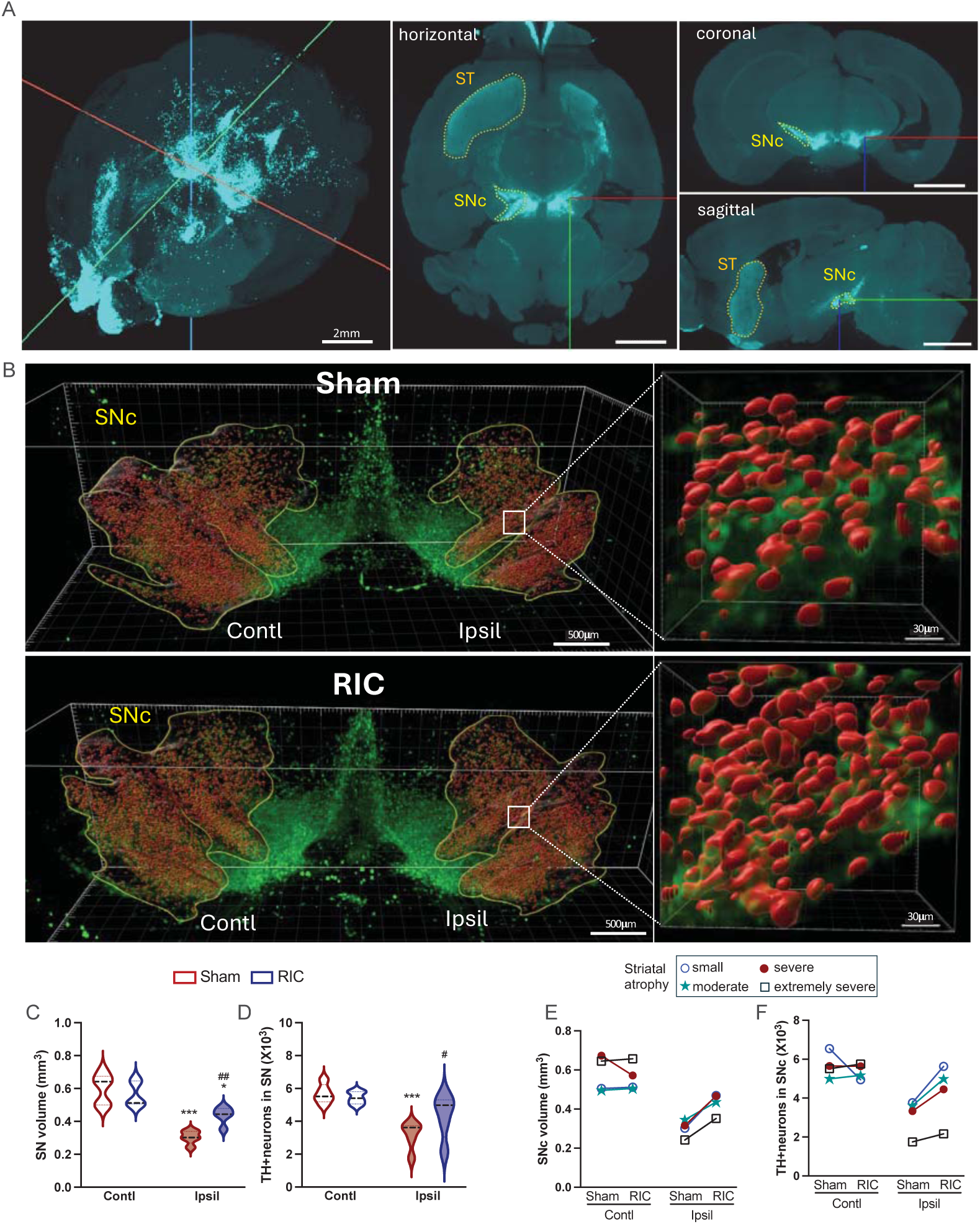
RIC attenuates dopaminergic neuron loss in substantia nigra pars compacta (SNc) in chronic stroke. Mice expressed GFP in tyrosine hydroxylase (TH) neuron (Tg^TH-GFP+^) were subjected 30 min MCAO and sham or RIC were applied 2 h after reperfusion. Brain images were obtained at 2-month post-ischemia. N=5/group. **A**, 3D whole brain image with horizontal, coronal, and sagittal views. GFP+ expression in TH+ cells. ST, striatum; SNc, substantia nigra compacta. **B**, 3D images of SNc indicated by yellow boundaries and the red represents areas where TH+ neurons are present. High resolution of SNc showing individual TH+ neurons in the ipsilateral hemisphere. **C, D,** Quantification of SNc volume (indicated by yellow outline) and TH+ neurons indicated by red in yellow boundaries. **E, F** Pairwise analysis of SNc volume (C), and TH+ neuron number (D) based on the severity of striatal atrophy. Contl, contralateral; Ipsil, ipsilateral; Sham, Sham conditioning; RIC, Remote ischemic conditioning, Two-way ANOVA followed by *post hoc* Fisher’s LSD test. *** *p* < 0.001 Contl vs. Ipsil (Effect of stroke), ##, ### *p* < 0.01, 0.001 Sham vs. RIC (Effect of RIC).

To address whether RIC-rescued SNc volume and TH+ neurons is the result of reduced striatal infarction in RIC animals, we compared animals based on the severity of striatal atrophy. At 2 months post-stroke, striatal atrophy varied within both sham and RIC groups, but the mean striatal atrophy was comparable between the groups (Fig S11). Pairwise comparison between sham and RIC groups in animals with small, moderate, or large atrophy showed that RIC mice displayed larger SNc volumes (Figure 7E) and higher TH+ neuronal cell counts (Figure 7F) in the ipsilateral SN when compared to sham animal with similar atrophy. The striatal atrophy-independent rescue of SNc structure suggests that RIC limits transneuronal degeneration independent of stroke-induced primary infarction.

### Mφ-CD36 deletion abrogates RIC-attenuated transneuronal degeneration and enhances stroke recovery

To investigate the role of CD36 in RIC-induced structure and function relationship, TH+ neuron loss in SNc and functional recovery were assessed in WT and cKO^MMϕ^ mice. Compared to sham mice, RIC significantly increased the number of surviving TH+ neurons in the ipsilateral hemisphere at 2m post-stroke (Figure 8A). The attenuation of the TH+ cell loss is associated with improved motor function in the open field, increased grip strength, and rotarod performance in chronic stroke (Figure 8C-F). In cKO^MMϕ^, RIC did not attenuate TH+ neuron loss (Figure 8B). The absence of protection from secondary degeneration is associated with no improvement in motor function (Figure 8G, H, J). and grip strength, which was significantly worse in RIC mice (Figure 8I). The structural and functional studies thus showed that CD36 mediates RIC-attenuated transneuronal degeneration in SNc and stroke recovery. These results collectively showed the importance of CD36 in the RIC effect by upregulation of CD36 expression in circulating monocytes, increased their entry into the injured brain, enhanced efferocytosis during an acute phase of stroke, mitigating transneuronal degeneration, and promoting functional recovery in chronic stroke (Figure 8K).

**Figure 8.**
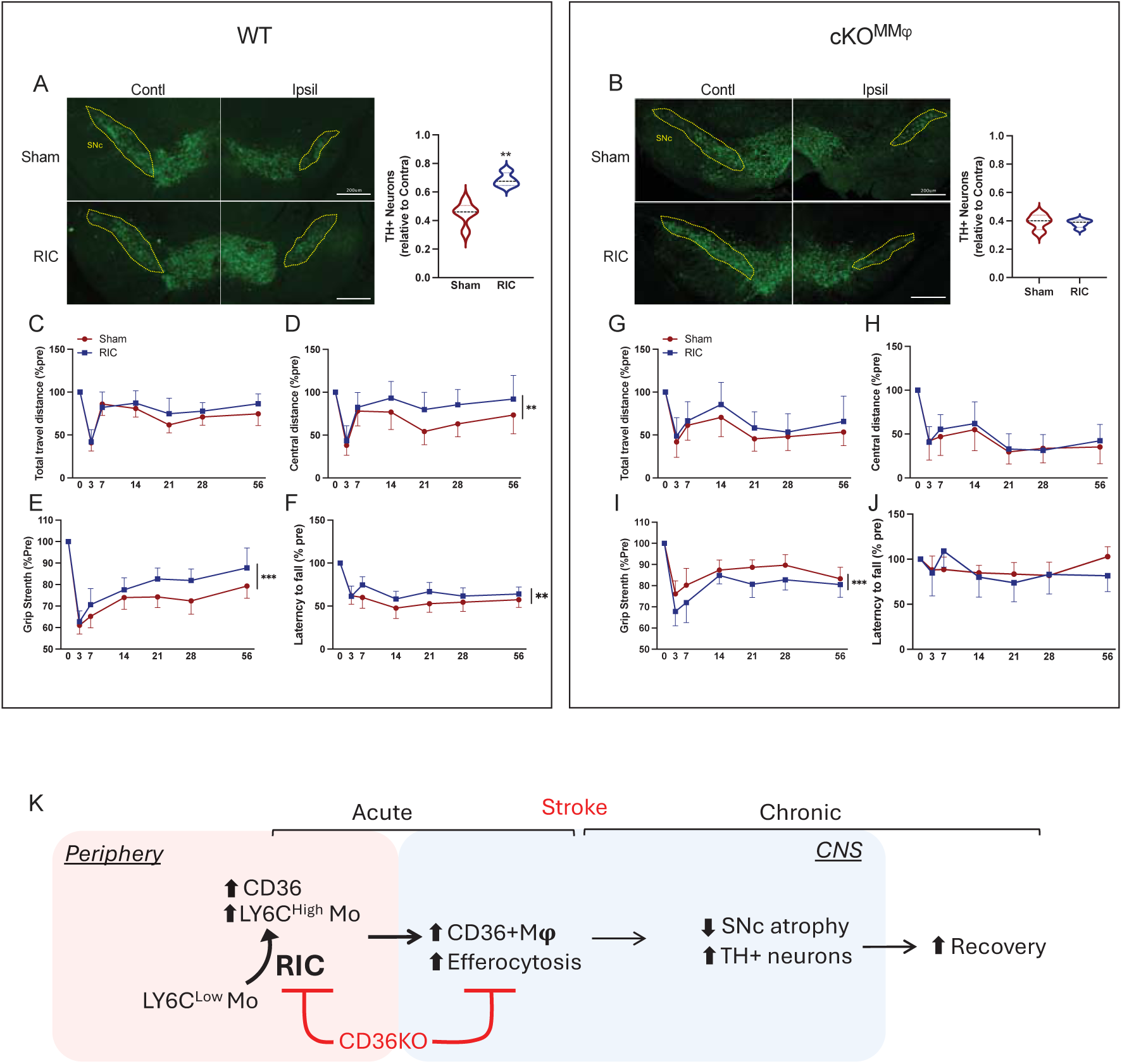
Mφ-CD36 deletion abrogates RIC-attenuated transneuronal degeneration and enhances stroke recovery. WT (CD36^fl/fl^) and cKO^MMφ^ mice with MCAO were received sham or RIC at 2h after MCAO. N=4-5/group/genotype. **A, B,** TH immunofluorescence staining in the brain sections at 2 months after MCAO in WT (A) and cKO^MMφ^ (B) mice. The number of TH+ cells was measured in both hemispheres and expressed as ratios of Ipsilateral/Contralateral. T-test: ** *p*<0.001. **C-F**, Longitudinal post-stroke behavior test in WT mice (n=28 sham, 19 RIC), **G-J** Longitudinal post-stroke behavior test cKO^MMφ^ mice, n=14/group. Open field test to assess total travel distance and distance in the center zone (C, D, G, H). Grip strength in the hind limbs (E, I) and rotarod (F, J). Data were presented as percent of pre-stroke baseline (% pre), Two-way ANOVA: *, **, *** *p* < 0.05, 0.01, 0.001 WT vs. cKO^MMφ^ (effect of genotype). K. Diagram depicting RIC-induced biological responses in stroke.

## Discussion

This study demonstrates that CD36-mediated efferocytosis in monocyte-derived Mφ is a critical mechanistic component of RIC that mediates the attenuation of transneuronal degeneration and promotes stroke recovery. Neural inflammation followed by the resolution of inflammation are cardinal features of stroke pathophysiology. These processes are dependent on infiltration of peripheral immune cells, primarily monocytes, into the injured brain. These pivotal roles played by monocytes advocates for the development of strategies to target these cells in order to influence injury progression and tissue repair. RIC is as a promising noninvasive and clinically viable approach for cross-organ protection ^9,10^, but the application of RIC in stroke patients has been controversial. The current study, however, provides novel insight into the molecular and cellular events underlying the acute changes in the blood following RIC that facilitate long-term functional recovery in experimental cerebral ischemia. A key finding in this study is that RIC-enhanced CD36 expression in monocytes and CD36-mediated efferocytosis in Mφs is required to attenuate delayed transneuronal degeneration in the post-ischemic brain and promote stroke recovery. Together, these findings suggest that RIC is an effective strategy to manipulate monocytes in the peripheral immune system that influence injury progression and tissue repair following stroke.

### The mechanism by which RIC increases CD36 expression in the periphery

An important event triggered by RIC is an increase in CD36 MFI within circulating monocytes that does not alter the total number of CD36+ monocytes (Figure 3C, D). A likely mechanism for the increase in CD36 expression within monocytes is the availability of CD36 ligands in circulation. Repeated episodes of ischemia-reperfusion with RIC produce oxidative stress and a sudden release of reactive oxygen/nitrogen species from mitochondria ^59–61^. Activation of CD36 by the presence of abundant ligands (e.g., oxidized or modified low-density lipoproteins Ox/m LDLs) generated by RIC drive a feed-forward upregulation of CD36 expression ^62,63^. An important finding in this study is that the increase in CD36 expression is specific to Ly6C^High^ monocytes. The absence of CD36 upregulation in Ly6C^Low^ monocytes could be due to the higher CD36 baseline in this subset (Figure. 3F), which may reflect the patrolling nature of this subset along the endothelium for interacting damaged and apoptotic cells, and modified forms of LDL in the vasculature ^64^. Ly6C^High^ monocytes, however, are the primary monocyte that infiltrates the injured brain via the CCR2-MCP1 axis, and their migration underlies the elevated levels of CD36+/CD45^High^ Mφ in the post-ischemic brain (Figure. 2E, F).

### CD36-mediated efferocytosis in the injured CNS

The function of CD36 in eliciting and suppressing inflammation is context-dependent ^37,65,66^. Given that CD36 functions in clearing cellular debris for tissue repair, the RIC-driven enhancement of CD45^High^ Mφ with greater efferocytosis activity is the cellular mechanisms that underlies the protective effect of RIC in the post-ischemic brain. An important question, however, is how the enhanced efferocytosis in infarcted tissue during acute stroke preserves TH+ neurons at 2 months post-stroke and promotes long-term stroke recovery in a CD36-dependent manner. Efferocytosis is a crucial step in resolving tissue injury that converts inflammatory Mφs to a reparative phenotype ^6^ that is proliferating and pro-resolving ^33^. *In situ* tracking of labeled monocytes in mice with stroke showed that monocyte-derived Mφs are the predominant phagocytes that also have significantly greater efferocytosis capacity than microglia in the ischemic brain ^6^. The ability of RIC to promote resolution of inflammation in stroke is supported by studies that have reported a reduction both in perihematomal edema in patients with hemorrhagic stroke ^67^ and brain swelling in mice with ischemic stroke ^2^. Mechanistically, this resolution requires CD36, as evidenced by lowered efferocytosis levels when CD36 is deleted in Mφs, but the lack of CD36 does not completely abolish efferocytosis (Figure 5) and indicates that there are additional receptors that engage efferocytosis. Moreover, several factors also influence efferocytosis levels, including the nature of phagocytes (professional vs. non-professional), and the ratio between phagocytic and apoptotic cells ^68,69^. In addition, efferocytosis can occur in either single or multiple, continuous rounds, the latter of which may be CD36-dependent. Continual efferocytosis is mechanistically distinct from a single round of efferocytosis and relies heavily on the metabolism of ingested apoptotic cell cargo ^46^. Disruption of continual efferocytosis delays injury resolution and expands the necrotic core in atherosclerosis ^70^. Transporting long-chain fatty acids via CD36 to the cells provides fuels for fatty acid oxidation, which sustains the activation of reparative Mφs ^66^, and may facilitate RIC-enhanced continual efferocytosis and tissue resolution.

### CD36 in RIC-rescued structure and function

Proximal MCA occlusion produces an infarct that is primarily in the striatum, but also part of the cortex. The long-term injury, however, is not restricted to the primary injury sites because damage spreads into remote areas in a delayed manner, such as the thalamus and SNc ^7^. In this study, the structure-function analyses demonstrated that RIC preserves SNc neurons and enhances stroke recovery in a CD36-dependent manner. The faster, CD36-dependent resolution in the acute phase of stroke likely accounts for the protection of the secondary structure and functional benefit in RIC animals. Notably, RIC rescue of SNc volume and TH+ neurons in the ipsilateral hemisphere was not dependent of the severity of striatal atrophies (except for one animal with extremely severe striatal atrophy; Figure 7E, F). Similarly, RIC-enhanced motor recovery in mice with severe stroke was also comparable to the those with smaller infarction ^2^. Therefore, the CD36-dependent, but infarct size-independent, benefits of RIC indicate that a timely and rapid resolution, rather than infarct severity, in the primary injury sites determines subsequent structure integrity and behavior. This also suggests that CD36 is a key target for accelerating injury resolution with RIC-induced structural and functional benefits.

### Peripheral markers to predict RIC benefit

Stroke with brain atrophy is associated with less favorable recovery ^71^, and retrospective imaging studies in ischemic stroke patients have shown secondary degeneration in the SN is associated with poor stroke outcomes. ^72,73^. The findings in this study show that RIC can attenuate secondary degeneration and promote recovery, which suggests that RIC can be a clinical strategy to preserve brain structural integrity in remote areas and improve stroke recovery. The outcome of RIC clinical trials on acute stroke patients has been controversial ^24–26^, likely due to several issues, including inconsistency include age, comorbidities, stroke subtypes, the time of RIC initiation, and compliance ^74–78^. By defining important mechanistic events underlying RIC, as we have done in this study, this will facilitate the establishment of mechanism-based protocols for RIC dosage, frequency, and intervals in repeated RIC applications. Using RIC to target peripheral changes in CD36+ monocytes, this non-CNS-directed immune strategy may also enable optimization of RIC protocols to use minimal dosing at maximal intervals to improve compliance issues.

### Limitations

Previous studies have demonstrated phenotypic plasticity of phagocytes in the injured tissue environment and efferocytosis can drive peripheral-derived CD45^High^ Mφs to adopt a “microglia-like” CD45^Low^ phenotype ^6,7^. These phenotype changes in monocyte-derived Mφs prevented assessing the contribution of CD45^Low^ Mφ to efferocytosis because the CD45^Low^ subset also contains resident microglia. Since efferocytosis can promote proliferation of pro-resolving Mφs ^33^, we suspect that the increased bead+ cells in the CD45^Low^ subset (Figure 1H) likely due to the proliferation of efferocytosing Mφs that have changed to a reparative Mφ phenotype. Another limitation is the impact of co-morbidities on RIC benefits. Recent studies indicate that obesity impedes the RIC-mediated increase in CD36 expression and monocyte shift, and the disruption of these peripheral changes prevents improvements in recovery in stroke ^78^. Further studies are required to assess how RIC interacts with other co-morbidities, including obesity, diabetes, and aging, to optimize RIC-based interventions and improve stroke recovery.

## Supporting information

Detailed methods are provided in the Supplemental material

Major Resources Table are provided in the Supplemental material

## Acknowledgments

The authors gratefully acknowledge the Burke Neurological Institute Structural and Functional Imaging Core for their support and assistance in this work. We also thank Dr. Yutaka Yoshida for providing access to Imaris software.

## Author contributions

HJ performed efferocytosis assay and flow cytometry analyses in vivo or in vitro studies and wrote the manuscript. IK performed stroke modeling. IP performed immunofluorescence studies, and cell counts and volumetric measurements in 3D images. SM processed the 3D volumetric data. KP performed immunofluorescence studies. JM performed the neurological assessment. AM preformed brain atrophy measurement. WW performed whole brain clearing, labeling, and imaging. XW generated the 3D visualization. ZW designed the whole mount test together with collaborators. JY generated cKO^MMφ^ mice line and established protocols for RIC and flow analyses. MF generated CD36^fl/fl^ mice and wrote the manuscript. JC provided TH-GFP mice and wrote the manuscript. SC designed the overall study, analyzed data, and wrote the manuscript.

## Non-standard Abbreviations and Acronyms

Acs: apoptotic cells
CCR2: C-C motif chemokine receptor 2
cKO^MMφ^: mice with a specific deletion of CD36 in monocytes/Mφs
Contl: Contralateral hemisphere
Ipsil: Ipsilateral hemisphere
MCAO: Middle cerebral artery occlusion
Mo: monocytes
Mφ: macrophages
RIC: Remote ischemic limb conditioning
SNc: Substantia nigra pars compacta
ST: Striatum
TH: tyrosine hydroxylase
WT: Wild type

---

**Figure S1.**
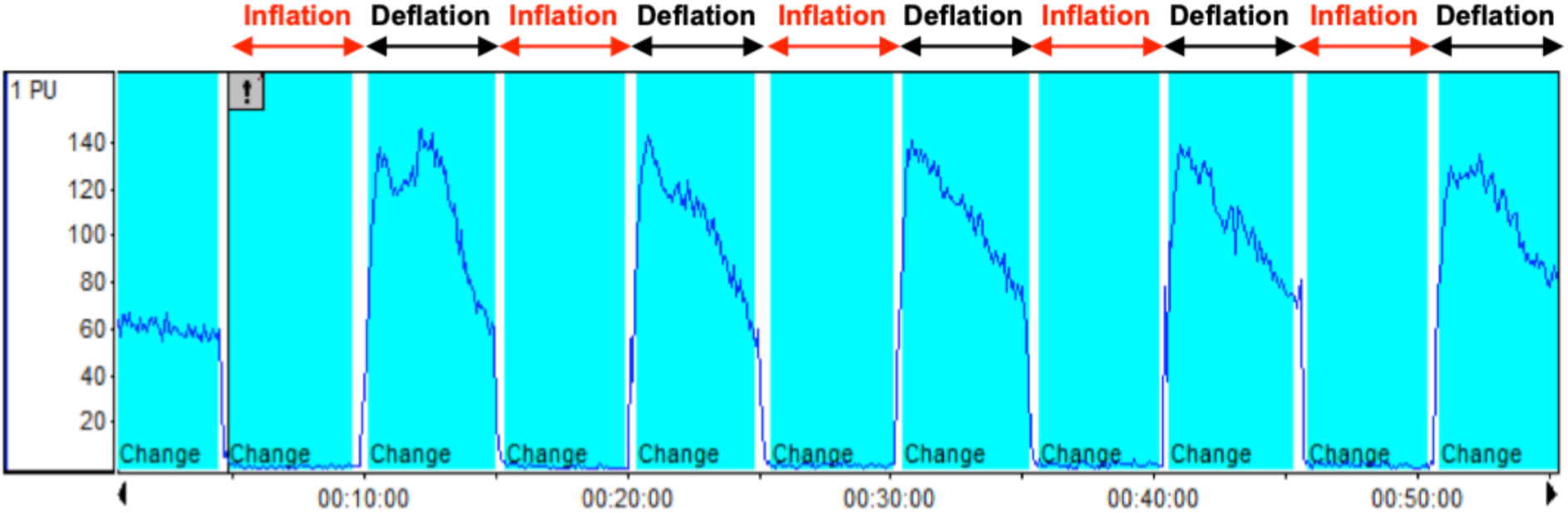
Remote limb ischemic conditioning (RIC) RIC was performed on the left hindlimb of the mice by applying a total of 5 cycles of inflation and deflation (5 min X 5 min intervals between cycles). The hindlimb blood flow was measured using Laser-Doppler flowmetry.

**Figure S2.**
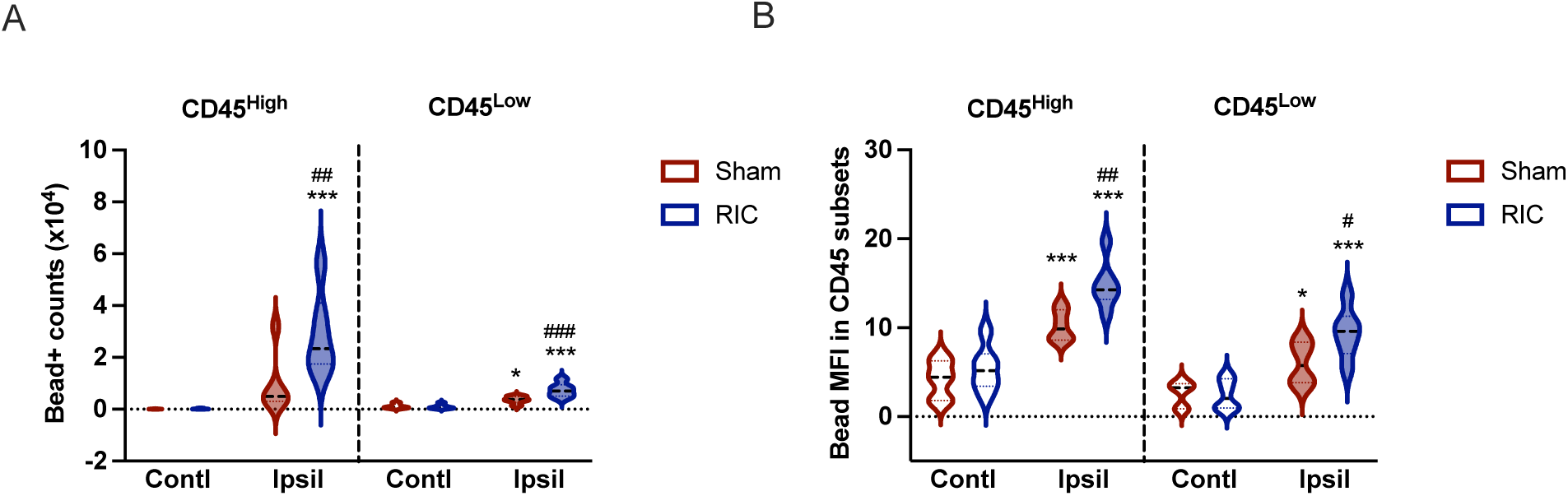
*In vivo* efferocytosis assay. The effect of RIC on efferocytosis in CD45 subsets in the brain immune cells isolated from 3d post-ischemia. Brain immune cells were gated for [CD45+/CD11b+/Lin-] Mφ/microglia and assessed for number (**A**) and mean fluorescent intensity (MFI) (**B**) of bead+ cells in CD45^High^ and CD45^Low^ subsets. N=6/group. Statistical significance was assessed with two-way ANOVA followed by post hoc Fisher’s LSD test. ^*,^ *** *p* < 0.05, 0.001 Contl vs. Ipsil (Effect of stroke); ^#, ##, ###^ *p* < 0.05, 0.01, 0.001 Sham vs. RIC (Effect of RIC).

**Figure S3.**
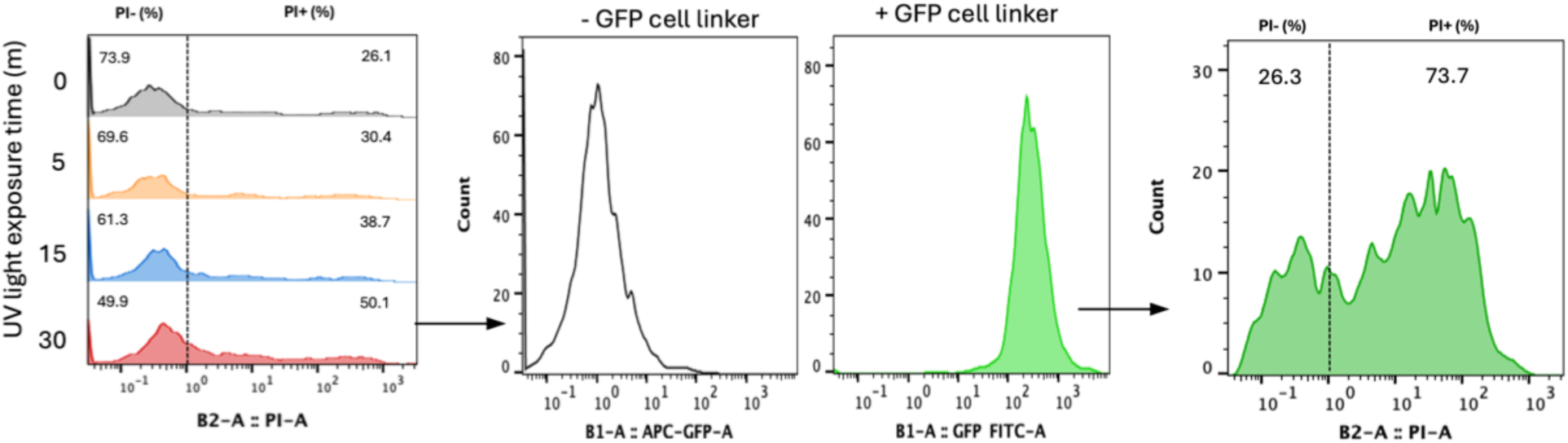
Generation of apoptotic cells (ACs). Isolated splenocytes incubated under UV light for 30 min cause the half of the cells to become apoptotic (propidium Iodide PI+). Incubation of cells with GFP Fluorescent Cell Linker labels the lipid region of the cell membrane (>99% efficiency). Approximately 70% GFP+ cells are PI+ (AC^GFP^).

**Figure S4.**
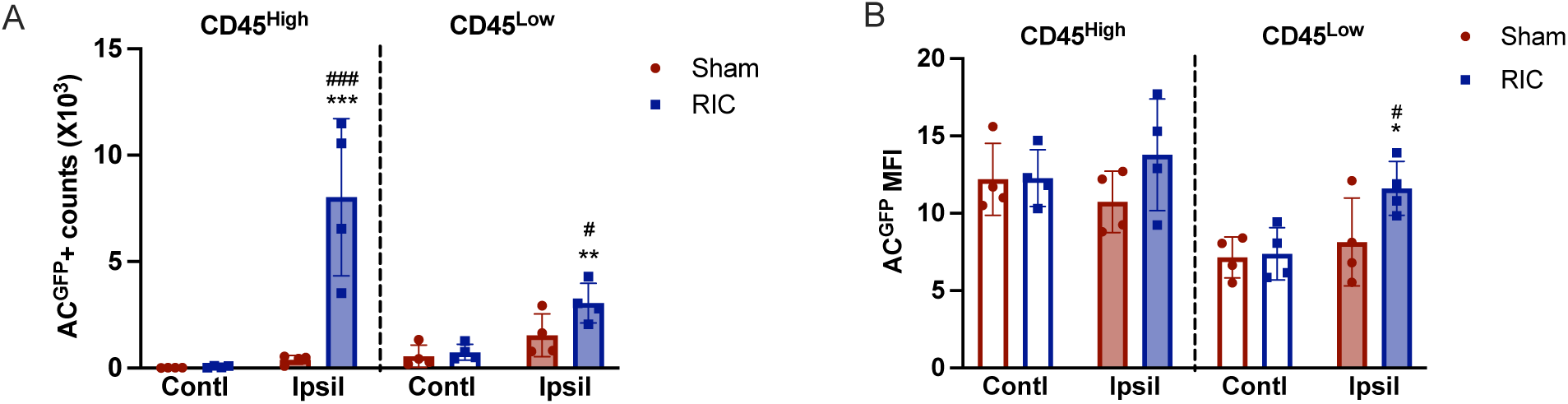
*In vitro* efferocytosis assay. The effect of RIC on efferocytosis in CD45 subsets assessed in the brain immune cells isolated from 3d post-ischemia. Brain immune cells were gated for Mφ/microglia [CD45+/CD11b+/Lin-] and assessed for number (**A**) and mean fluorescent intensity (MFI) (**B**) of AC^GFP^+ cells in CD45^High^ and CD45^Low^ subsets. N=4/group. Statistical significance was assessed with two-way ANOVA followed by *post hoc* Fisher’s LSD test. ^*, **,***^ *p* < 0.05,0.01, 0.001 Contl vs. Ipsil (Effect of stroke); ^#, ###^ *p* < 0.05, 0.001 Sham vs. RIC (Effect of RIC).

**Figure S5.**
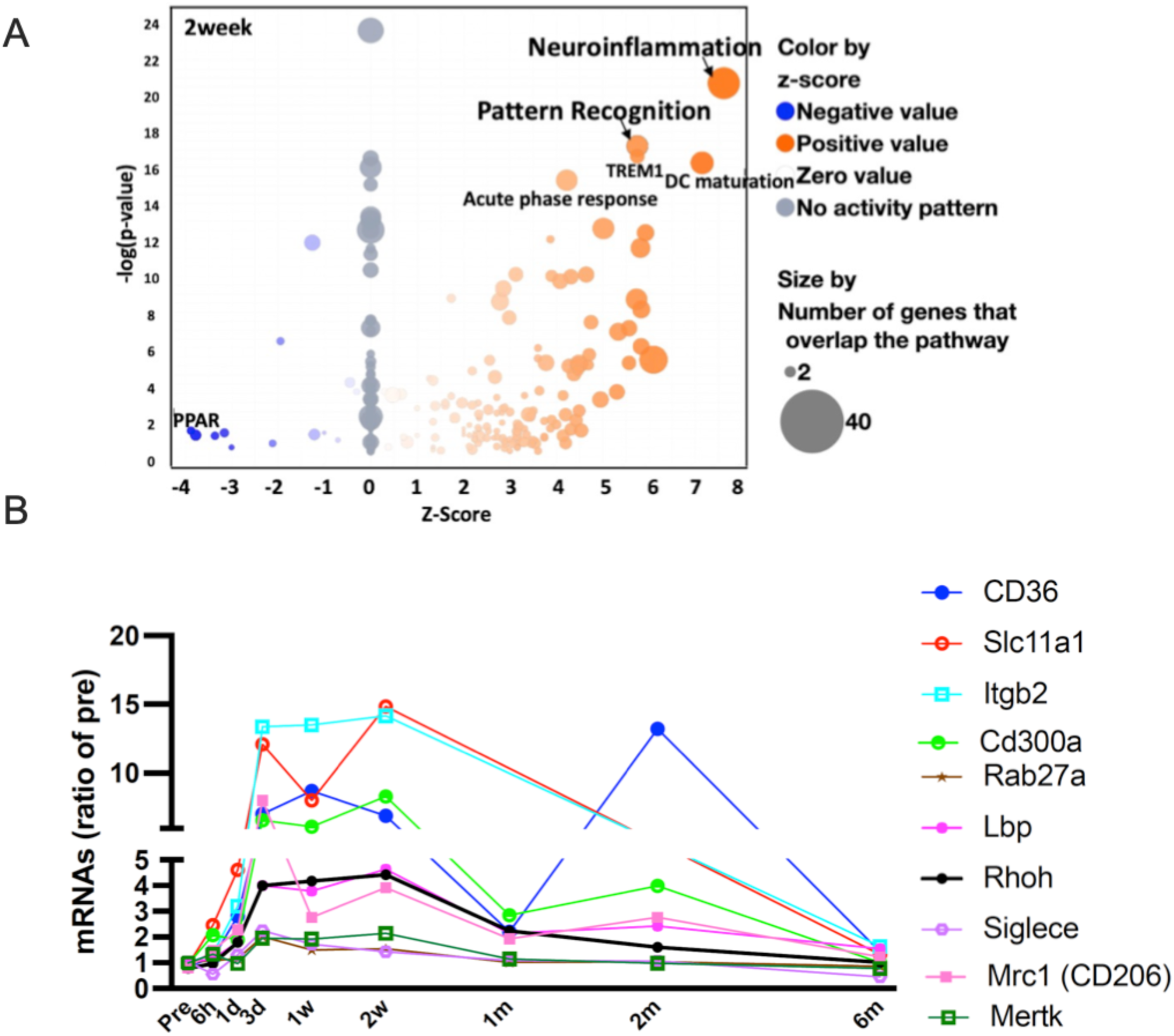
Stroke persistently upregulates the genes pertaining to pattern recognition. **A,** Ingenuity canonical pathway analysis. Volcano plot of RNA-seq data at 2w after stroke showed the most upregulated gene signatures are neuroinflammation and pattern recognition. **B**, Sustained upregulation of genes in the ipsilateral hemisphere from an acute phase out to the 6m recovery phase of stroke are related to pattern recognitions.

**Figure S6.**
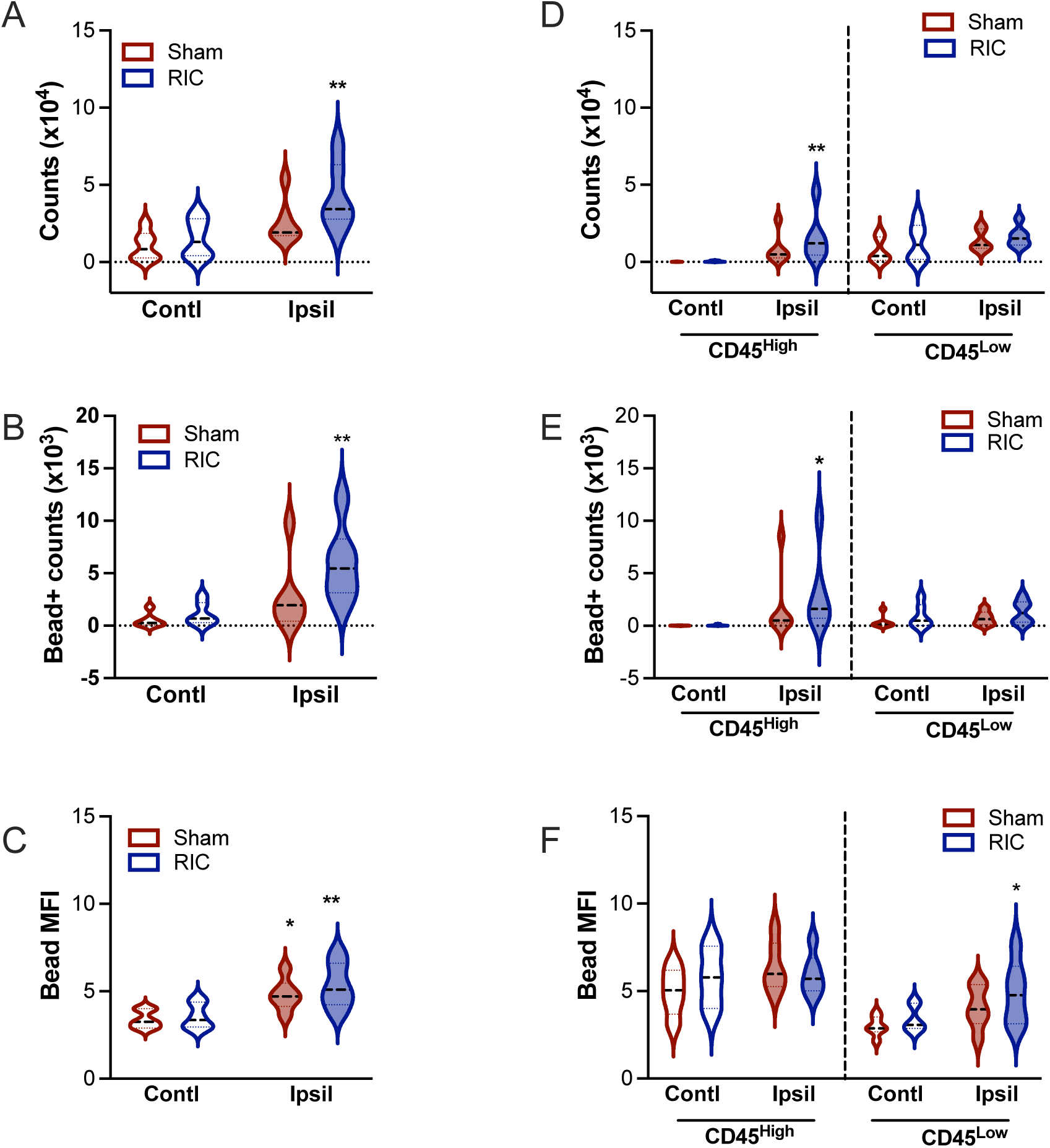
RIC-enhanced efferocytosis does not occur in cells that do not express CD36. All assessments were performed in [CD45+/CD11b+/Lin-] populations **A**, Quantification of the number of CD36-negative cells. **B**, the number of bead+/CD36-cells, **C,** Bead MFI in CD36-cells, **D**, Quantification of the number of CD36-negative cells in CD45^High^ and CD45^Low^ subsets. **E,** the number of bead+/CD36-cells in CD45 subsets, **F,** Bead MFI in CD36-cells. N=6/group. Statistical significance was assessed with two-way ANOVA followed by *post hoc* Fisher’s LSD test. ^*,^ \*\**p* < 0.05, 0.01, Contl vs. Ipsil (Effect of stroke); no RIC effects are observed. Abbreviations: Contl (contralateral); Ipsil (ipsilateral); Sham (sham conditioning); RIC (remote ischemic conditioning).

**Figure S7.**
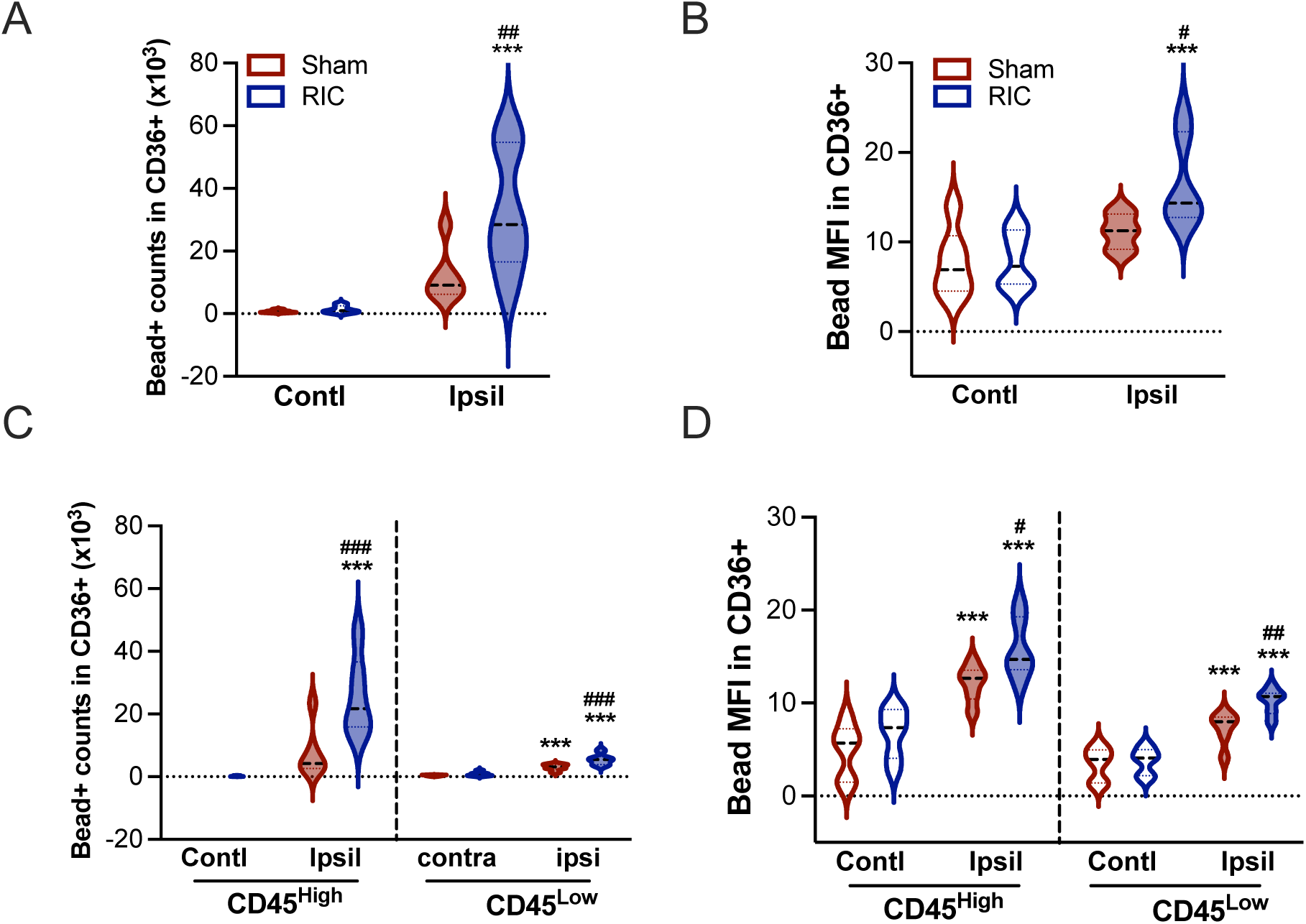
RIC increased the number of beads+ cells and bead intensity in cells that express CD36. Efferocytosis assays were performed in [CD45+/CD11b+/Lin-] populations. **A, B**, Quantifications of the number of bead+ (A) and bead intensity (B) in CD36+ cells. **C, D**, Quantifications of the number of bead+ cells (C) and bead intensity (D) in CD45^High^ and CD45^Low^ subsets. n=6/group. Contl, Contralateral; Ipsil, Ipsilateral; Sham, Sham conditioning; RIC, Remote ischemic conditioning. Statistical significance was assessed with two-way ANOVA followed by *post hoc* Fisher’s LSD test. *** *p* < 0.001 Contl vs. Ipsil (Effect of stroke); ^#, ##, ###^ *p* < 0.05, 0.01, 0.001 Sham vs. RIC (Effect of RIC).

**Figure S8.**
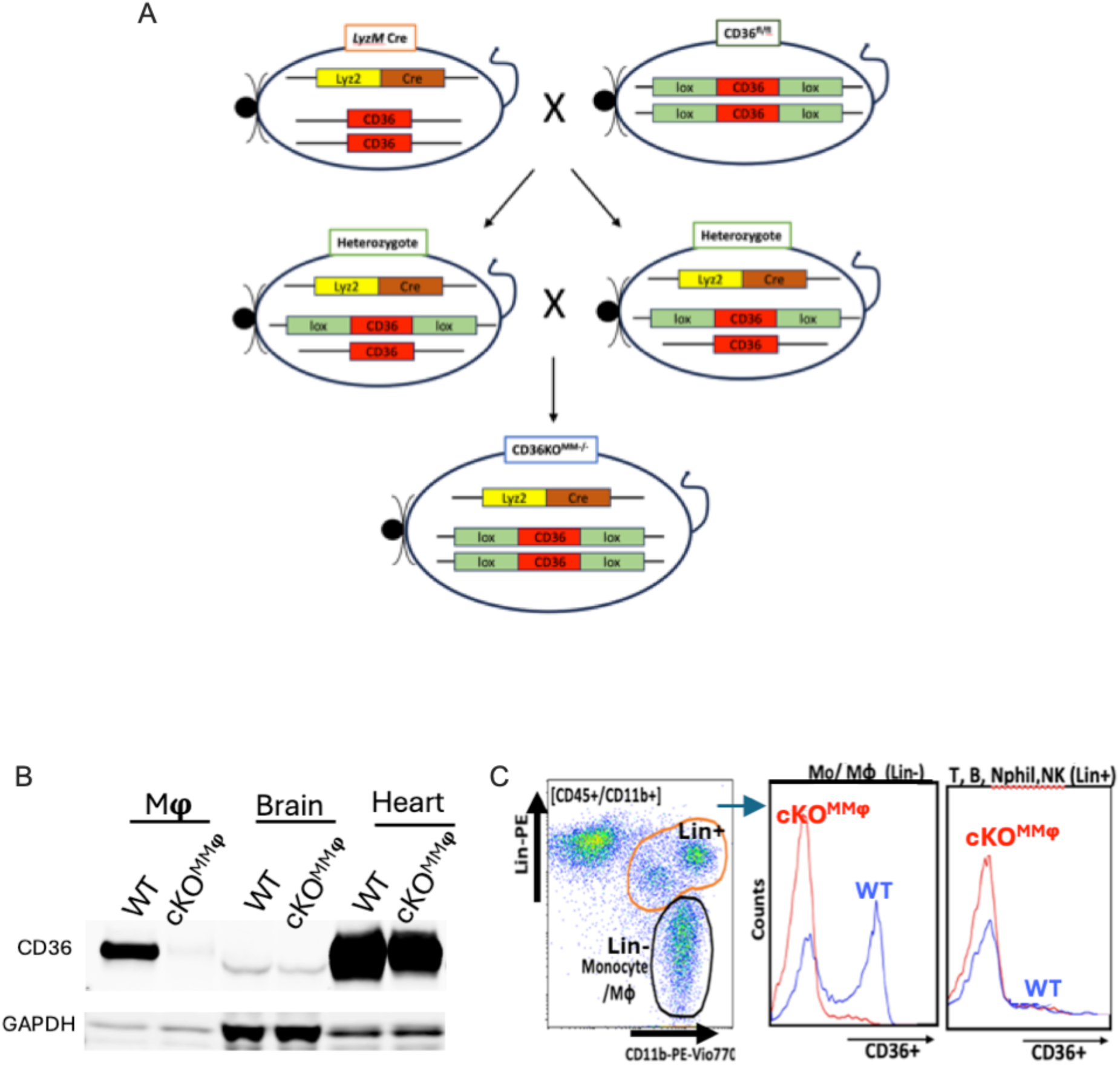
Generation and characterization of cKO^MMφ mice.^ **A,** Generation of cKO^MMφ^ mice. LoxP flanked CD36 in C57 background (CD36^fl/fl^, WT)^79^ were crossed with *LyzM* Cre mice (B6.129P2-*Lyz2^tm1(cre)Ifo^*, Jackson Lab) to generate mice with a conditional deletion of CD36 in monocytes/Mφ **B**, Characterization of cKO^MMφ^ mice. CD36 Western blots in CD36^fl/fl^ (WT) and cKO^MMφ^ mice. Note that cKO^MMφ^showed a selective CD36 deficiency in Mφ while retaining the expression in the brain (e.g., microvascular endothelial cells) and heart (cardiomyocytes). **C**, Flow cytometry analysis for CD36 expression in the blood. [Lin-] cells in the [CD45+/CD11b+] population for monocyte gating and [Lin+] for T- and B-cells, neutrophils, NK cells. Note that both WT (CD36^fl/fl^) and cKO^MMφ^ mice display extremely low levels of CD36 expression in T and B lymphocytes, NK cells, and neutrophils with no detectable differences between the genotypes.

**Figure S9.**
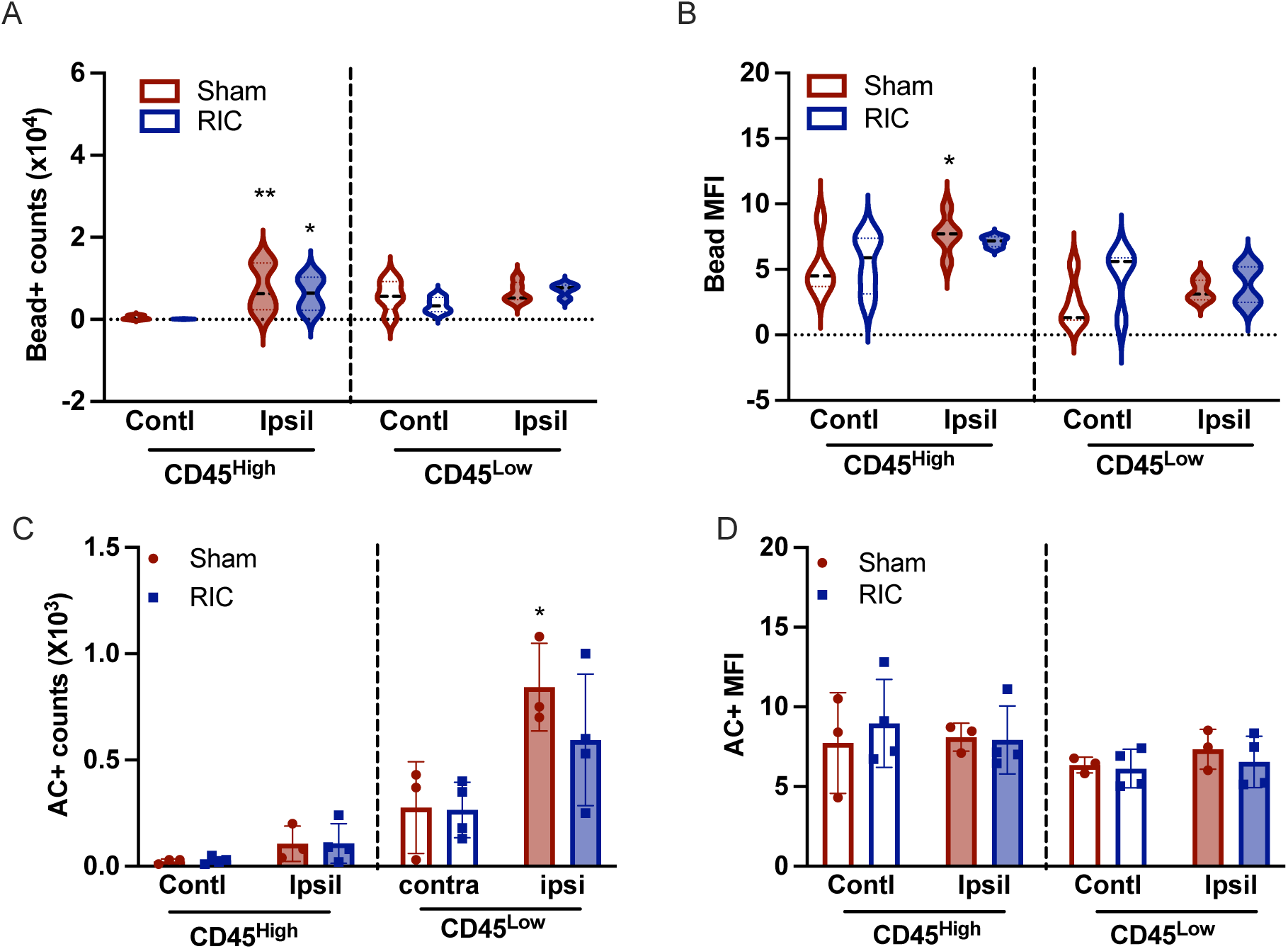
CD36 mediates RIC-enhanced efferocytosis in the post-ischemic brain. cKO^MMφ^ mice with 30 min MCAO were subjected to Sham or RIC at 2h post-MCAO. **A, B** *In vivo* efferocytosis assay, Number of Beads^580/605^+ cells (A) and MFI (B) in CD45^High^ and CD45^Low^ Mφs/microglia. **C, D,** *In vitro* efferocytosis assay, Number of apoptotic cell-containing (AC+) cells (C) and MFI of AC+ cells (D) in CD45^High^ and CD45^Low^ Mφs/microglia in cKO^MMφ^ mice. Contralateral; Ipsil, Ipsilateral; Sham, Sham conditioning; RIC, Remote ischemic conditioning. Statistical significance was assessed with two-way ANOVA followed by *post hoc* Fisher’s LSD test. ^*,^ ** *p* < 0.05, 0.01, Contl vs. Ipsil (Effect of stroke).

**Figure S10.**
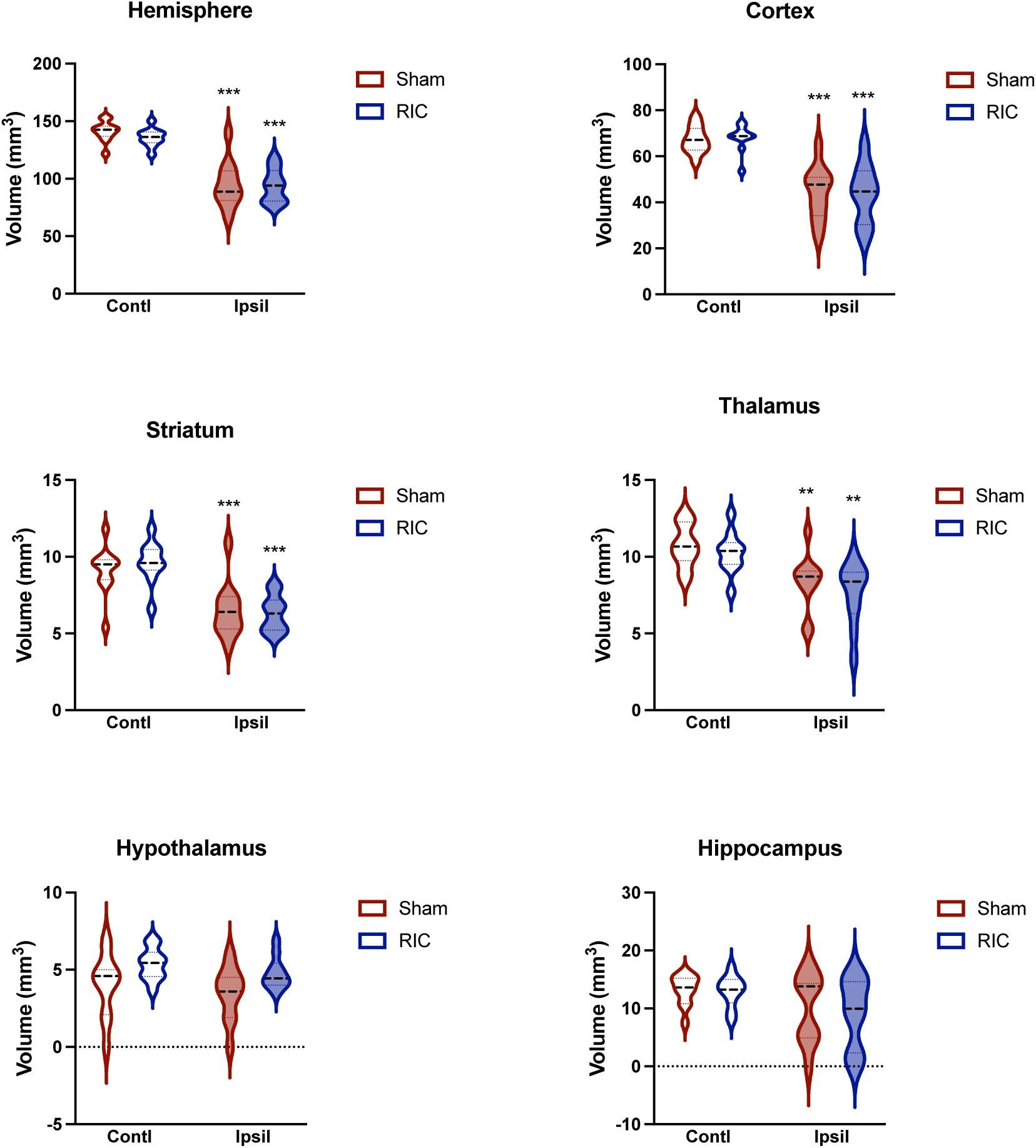
Assessment of atrophy in brain sub-regions at 2m post-stroke. C57 mice were subjected to MCAO and received sham or RIC 2h after MCAO. Volume assessment in the hemisphere, primary injury sites (striatum and cortex), and remote areas (thalamus. hypothalamus and hippocampus). n=10-11/group. Two-way ANOVA: **, *** *p* < 0.001, 0.001 Contl vs. Ipsil (Effect of stroke).

**Figure S11.**
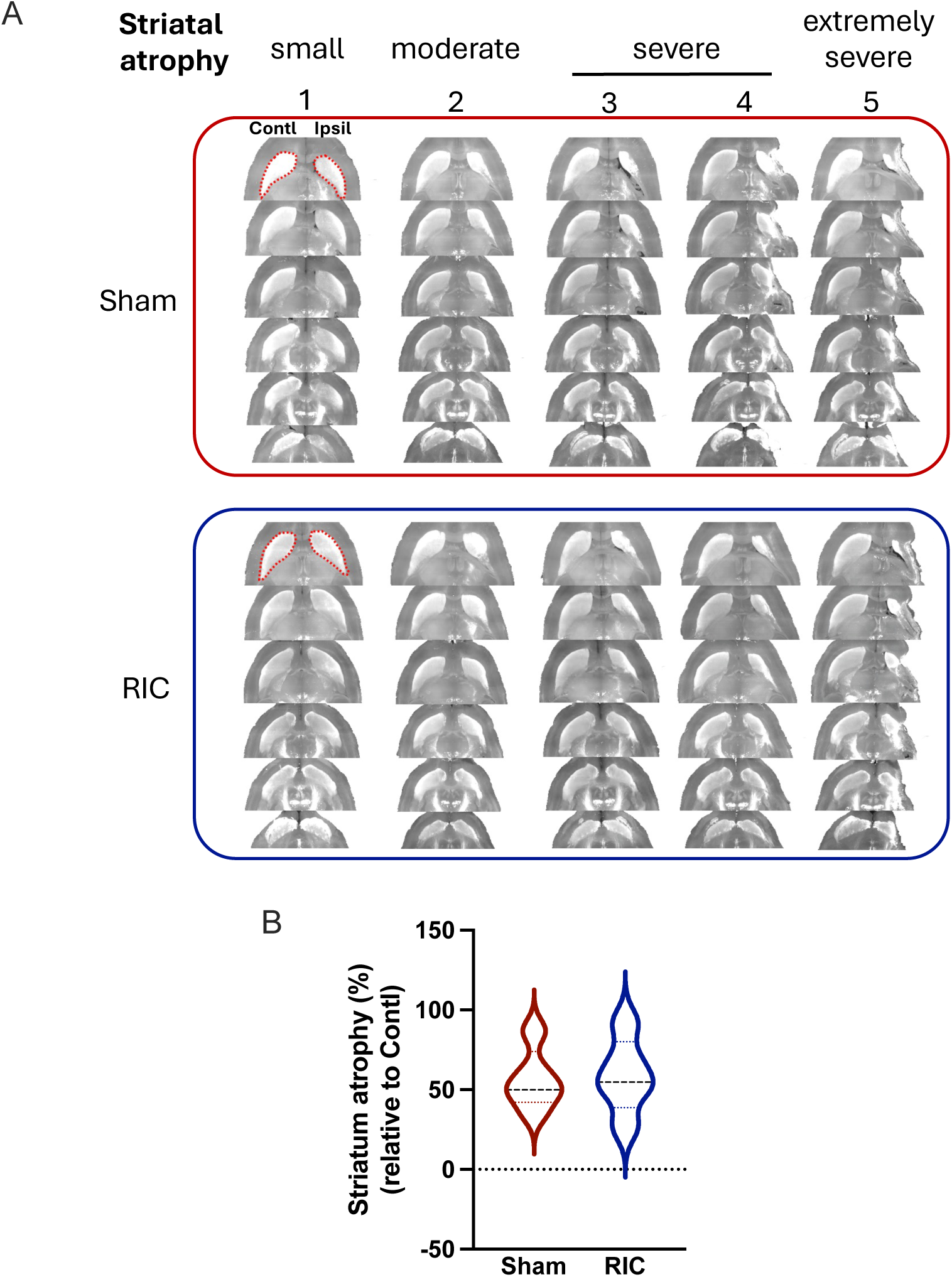
Effect of RIC on striatal injury size. **A,** Horizontal view of serial section images containing the striatum at 2 month post-stroke. Images were captured at 400 um intervals. **B,** Tissue atrophy in the ipsilateral striatum was expressed as percentages of the contralateral striatum. N=5/group.

