## Supplementary material for "Remote ischemic conditioning attenuates transneuronal degeneration and promotes stroke recovery via CD36-mediated efferocytosis": Detailed methods are provided in the Supplemental material

**Supplemental Methods**

**Animals:** The procedures for animal use were approved by the Institutional Animal Care and Use Committee (IACUC) of Weill Cornell Medical College. They were carried out in accordance with guidelines from the National Institutes of Health and Animal Research: Reporting of In Vivo Experiments (ARRIVE). The animals were maintained under controlled conditions with regulated temperature and humidity and on a 12 h light/dark cycle. Each cage contained a maximum of 5 mice and was equipped with ventilation and irradiated bedding (1/8-inch Bed O's Cobs, The Anderson, Maumee, OH). Sterilized food (PicoLab Rodent diet 5053, LabDiet, St. Louis, MO) and water were readily available *ad libitum* in the cages. Mice with CD36 conditionally knocked out in monocyte/Mφ ($cKO^{MM\varphi}$) were generated by crossing mice containing loxP flanked CD36 in C57 background ($CD36^{fl/fl}$; from Maria Febbraio) [79] with LyzM Cre mice (B6.129P2-Lyz2tm1(cre)Ifo; Jackson Laboratory). The $cKO^{MM\varphi}$ mice have a CD36 specifically deleted in macrophages without disrupting CD36 expression in other tissue (Figure S8). All experiments used male and female mice aged 3-4-month. TH-GFP+ mice (from Kazuto Kobayashi) expressed GFP in TH+ dopaminergic neurons [80] were used to investigate for transneuronal degeneration. These genetically modified mice are C57BL/6 congenic. All experiments used male and female mice aged 3-4-month.

**Transient middle cerebral artery occlusion (MCAO):** The procedure for 30-minute transient MCAO and post-stroke care has been described previously [81]. Briefly, mice were anesthetized with a mixture of isoflurane, oxygen, and nitrogen. A 6-0 Teflon-coated black monofilament surgical suture (Doccol, Redland, CA) was inserted into the exposed external carotid artery and advanced into the internal carotid artery until it was wedged into the Circle of Willis, obstructing the origin of the middle cerebral artery (MCA). The filament was left in place for 30 minutes before being removed. Cerebral blood flow (CBF) was measured before, during, and after the stroke using Laser-Doppler flowmetry (Periflux System 5010; Perimed, Järfälla, Sweden). Only animals that had both >80% reduction of pre-ischemic baseline CBF during MCAO and CBF >80% of baseline after 10 min of reperfusion were included in the study. Body temperature was maintained at 37 ± 0.5°C during, and for 30 minutes after, the MCA procedure using a rectal probe connected to a thermocouple-regulated heating water coil on the surgical board. Mice were placed in a recovery cage, and their body temperatures were maintained at 37 ± 0.5°C until they regained consciousness and resumed activity, after which they were returned to their home cages. During the acute phase, the mice were administered warm saline subcutaneously to prevent dehydration. The animals were given softened food and hydrogel (Clear $H_2O$) during the first week of recovery. Typically, the mice began to regain body weight at 3-5 days post-stroke and continued to recover.

**Remote limb ischemic conditioning (RIC):** RIC was performed on the left hindlimb of the isoflurane-anesthetized mice. This involved applying a total of 5 cycles of inflation and deflation (200 mmHg, 5 minutes each, with a 5-minute interval between cycles) using a small blood pressure cuff (Hokanson, Bellevue, WA). In a poststroke application, RIC was applied 2 h after reperfusion. Blood flow in the left hindlimb was measured during RIC and monitored by Laser-Doppler flowmetry (Periflux System 5010; Perimed, Järfälla, Sweden) (Figure S1). The Sham group, exposed to the same duration of isoflurane, served as a control.

**Assessment of monocyte subsets *in vitro*:** Sera were collected from Sham or RIC mice 24 h after MCAO surgery. The sera from each group (n=3 or 4) were combined and stored at -80°C until use. Naive splenocytes from WT or $cKO^{MM\varphi}$ were collected and prepared for single-cell suspensions. After the splenocyte collection, CD11b+ splenocytes were isolated using CD11b Microbeads (Miltenyi Biotec, San Diego, CA) following the manufacturer's instructions. The CD11b+ splenocytes were then incubated without serum, with Sham serum, or with RLC serum for 4 h at 37°C with 5% $CO_2$. After the 4h incubation, the cells were collected and incubated with antibody for flow cytometry. A total of 20,000 cells were analyzed.

**Brain immune cell isolation:** Mice were anesthetized with isoflurane and ketamine/xylazine before being perfused with ice-cold phosphate-buffered saline (PBS) containing heparin. Brains were removed, and the hemispheres were separated before being placed into ice-cold Hanks' balanced salt solution without $Ca^{2+}$ and $Mg^{2+}$ (HBSS, Life Technologies, Grand Island, NY). Tissue was enzymatically and mechanically dissociated using a MACS Neural Tissue Dissociation Kit (Miltenyi Biotec, San Diego, CA) and then treated with a myelin debris removal solution (Miltenyi Biotec, San Diego, CA). Isolated brain immune cells were used for either flow cytometer analyses or cultured for *in vitro* efferocytosis assays.

**Flow cytometry analysis:** From single-cell preparations, we determined the total cells by multiplying the dilution factors to the cell event reads. The primary antibodies used for cell staining were as follows: CD11b (PE-Vio770 anti-mouse CD11b REA, 1:50); CD45 (Vio-blue anti-mouse CD45 REA, 1:50); CD36 (Vio Bright B515 anti-mouse CD36 REA, 1:50) or (APC anti-mouse CD36 REA, 1:50); Ly6C (FITC anti-mouse Ly6C REA, 1:50) or (APC anti-mouse Ly6C REA, 1:50); and a mixture of lineage markers (Lin) conjugated with APC (APC-Lin) against T cells (APC anti-mouse CD90.2 REA), B cells (APC anti-mouse CD45R/B220 REA), natural killer cells (APC anti-mouse NK-1.1 REA, and APC anti-mouse CD49b REA), and granulocytes (APC anti-mouse Ly-6G REA) [29]. After a 20-minute incubation at room temperature in the dark, the cells were washed with PBS and then analyzed with a MACS Quant VYB flow cytometer (Miltenyi Biotec, San Diego, CA). The antibodies used for flow cytometry were purchased from Miltenyi Biotec. Antibody specificity was confirmed by analyzing cells only and using single and double antibody controls for validation.

**Efferocytosis assay:** For the *in vitro* assays, single immune cells isolated from the brain (2 x $10^5$/well) were cultured in 24-well plates in Macrophage-SFM medium (Thermo Fisher Scientific, Waltham, MA) containing 10% FBS for 2h at 37°C incubator. To simulate stroke milieu and to probe efferocytosis activity of the isolated brain immune cells, apoptotic cells (ACs) were co-cultured with the isolated immune cells. ACs were generated by placing splenocytes (1 x $10^7$ cells/100mm dish) under UV light for 30 min, which induces apoptosis (see Figure S3). The splenocytes were further labeled with GFP Fluorescent Cell Linker to label the lipid region of the cell membrane (MINI67/126, Sigma-Aldrich, St. Louis, MO), which allows >99% of cells to be labeled with the GFP, and ~70% of GFP+ splenocytes are ACs, based on propidium iodide staining. GFP+ apoptotic cells (1 X $10^6$) are added to the cultured immune cells (brain cells: APCs, 1:5) for 1h, washed with PBS. Co-cultures are harvested and analyzed by flow cytometry for efferocytosis of GFP+ cells in [CD45+ /CD11b+/Lin-] populations.

For *in vivo* assay, efferocytosis activity was determined by infusion of fluorescent microspheres (beads580/605; F-13083, Fisher Scientific, Waltham Thermo, MA) at 2d after MCAO via the retro-orbital venous sinus. Brain tissue was harvested one day later (3d post-

MCAO) and determined bead+ cells in [CD45+ /CD11b+/Lin-] populations by flow cytometry for the assessment of *in vivo* efferocytosis.

**Western blotting:** Brain tissue, heart, and peritoneal macrophages were homogenized and lysed in lysis buffer (C3228, Sigma-Aldrich, St. Louis, MO) with added complete Mini protease inhibitor (11836170001, Sigma-Aldrich). Lysates were centrifuged to obtain supernatants for protein analysis. Protein concentrations were determined, and equal amounts of protein were separated on NuPAGE 4–12% Bis-Tris gels (NP0322box, Thermo Fisher Scientific, Waltham, MA). After electrophoresis, proteins were transferred to PVDF membranes (#1620255, Bio-Rad, Hercules, CA). The membranes were blocked with Licor blocking buffer (#927-60001, Licorbio, Lincoln, NE) for 1 hour and then incubated overnight at 4°C with anti-mouse CD36 (1:2000, MAB1258, Sigma-Aldrich) or anti-rabbit GAPDH (1:5000, sc-25778, Santa Cruz Biotechnology, Dallas, TX) antibodies. After washing with Tris-buffered saline containing 0.05% Tween 20, membranes were incubated with IRDye 800CW donkey anti-mouse (1:2000, #926-32212, Licorbio) or IRDye 680CW goat anti-rabbit (1:2000, #926-68071, Licorbio) secondary antibodies in blocking buffer for 1 hour at room temperature. Protein bands were visualized using the Odyssey imaging system (Licorbio).

**Brain acute infarct size measurement:** Three days following MCAO, the brains were extracted, cryosectioned at a thickness of 20 μm, and collected at intervals of 600 μm. Infarct size and brain swelling were assessed using a total of 13 brain sections with Axiovision software (Carl Zeiss, Thornwood, NY), following the method outlined in a previous study [81]. The collected sections were examined under phase-contrast microscopy to identify the infarcted regions. The infarcted area was delineated in each section, and the infarct volume was calculated by multiplying the infarct area ($mm^2$) in each section by the distance to the next section (0.6 mm), then summing the values from all sections. The infarct volume was adjusted for edema by subtracting the difference between the hemispheric volumes. To determine the percentage of hemispheric swelling, the difference in volume between the ipsilateral and contralateral hemispheres was divided by the contralateral hemisphere volume and multiplied by 100.

**Whole brain imaging:** Fixed adult mouse brains from TH-GFP+ animals were delipidated using a modified Adipo-Clear protocol [82]. Briefly, brains were first washed with B1n buffer (H2O/0.1% Triton X-100/0.3 M glycine, pH 7), then transferred to a methanol gradient series (20%, 40%, 60%, 80% methanol in B1n buffer) using 4 mL for each brain and 1 hour for each step. This gradient series was followed by incubation in 100% methanol for 1 hour, then an overnight incubation in a 2:1 mixture of DCM:methanol and an 1.5-hour incubation in 100% dichloromethane (DCM) the following day. After which the brains were placed in 100% methanol for 1 hour three times, before treatment with a reverse methanol gradient series (80%, 60%, 40%, 20% methanol in B1n buffer) with 30 minutes for each step. Finally, brains were washed in B1n buffer for 1 hour, and then overnight in fresh B1n buffer. All these procedures were conducted at room temperature with rocking to ensure complete delipidation. The delipidated brain tissue was then blocked in PTxwH buffer (PBS/0.1% Triton X-100/0.05% Tween 20 containing 5% DMSO and 0.3M glycine) for 3 hours at 37°C before an overnight wash in fresh in PTxwH buffer at 37°C. Following this, tissue was successively washed at room temperature with fresh PTxwH buffer for 1 hour, 2 hours, and then overnight.

For staining, brain samples were immersed in a mixture of primary antibodies TH (Rabbit anti-TH, Thermo Fisher Scientific 701949, 1:100) and GFP (Mouse anti-GFP, Addgene 180084-rAb, 1:300) in PTxwH buffer for 14 days at 37°C. After which, samples were washed in PTxwH buffer for 1 h, 2 h, 4 h, and overnight, before three consecutive 1-day washes. Secondary antibodies used were AlexaFluor-594 (Jackson ImmunoResearch 711-587-003, donkey-anti-rabbit, 1:50) and AlexaFluor-647 (Jackson ImmunoResearch 115-607-186, goat-anti-mouse IgG2a, 1:250). Tissue was incubated with secondary antibodies were diluted in PTxwH for 10 days before being washed with PTxwH for 1 h, 2 h, 4 h, overnight, then three consecutive 1-day washes. The final wash was with PBS for one day. Tissues samples were then cleared with the iDISCO+ protocol [83]. Briefly, samples were dehydrated with a methanol gradient that started with water an concluded with 100% methanol, and followed by incubation with a DCM/methanol mixture overnight, and 100% DCM for 1 h twice the next day. Brain tissue was cleared for 4 h in dibenzyl ether and stored in a fresh dibenzyl ether before imaging.

Cleared brain samples were imaged horizontally in tiled sections with a LifeCanvas SmartSPIM lightsheet microscope. Imaging used 488/561/647 nm lasers with 3.6×/0.2 detection lens were used for green/red/far-red channels. Lightsheet illumination was focused with an NA 0.2 lense and axially scanned with an electrically tunable lense coupled to the camera (Hamamatsu Orca Fusion) in slit mode. The camera exposure was set at fast mode (2 ms) with 16-bit imaging with an X/Y sampling rate of 1.8 μm and Z step of 2 μm. Stitched lightsheet images were converted to a hierarchical chunked data organization (Neuroglancer Precomputed format) for visualization and processing for downstream analyses of regions of interest. Format conversion and hierarchical down sampling were performed using the Python library CloudVolume (version 8.24.2, https://github.com/seung-lab/cloud-volume/) and the Igneous software [84](version 4.20.3, https://github.com/seung-lab/igneous). Whole-brain 3D visualization was created with the ORS Dragonfly software using a downsampled version of the full image volume of the appropriate brain sample. Neuroglancer (https://github.com/google/neuroglancer) provides convenient 2D visualization of arbitrary orthogonal virtual cross-sections.

**Quantification of SNc volume and TH+ neuron count in the SNc region:**
3D image analysis used Imaris software version 10.1.1 (Bitplane AG, Zurich, Switzerland). Prior to analysis, the original whole brain stacks were flipped and cropped to provide smaller stacks that preserved the original voxel size, but only included regions encompassing the SNc. The same pre-processing procedure was used for each channel. Processed 3D image stacks were imported in Imaris at a voxel size of 1.8 x 1.8 x 2 μm and rendered using maximum intensity projections. Imported images were used to create a composite Imaris image object consisting of two channels with the first channel (pseudo-colored in green) containing the signal produced by TH+ neurons and the second channel (pseudo-colored in red) containing the signal produced by GFP expressing-TH+ neurons.

To isolate and quantify the neurons specifically within the SNc, freehand 3D regions of interest (ROI) around the left and right SNc volumes were created for each image object using manually drawn contours around the left and right SNc on every 55th 2D plane of the image object (corresponding to z-plane intervals of 100 μm). Autofluorescence signal in the green channel was used as an anatomical guide. Separate left and right SNc surfaces were created for each brain to enable independent quantification of the volume and the number of TH+ cells in the SNc of each hemisphere. The volume of each SNc region was calculated and provided by Imaris.

To count SNc TH+ cells, we developed a machine learning (ML)-based cell quantification procedure that utilized a pixel classifier provided by the ML Segmentation option in the automatic surface creation feature of Imaris. The pixel classifier was trained to segment TH+ cells using input from both channels and user's visual recognition. Training with both channels or with a cell having both bright cytoplasm and bright nuclei, rather than with only the green channel with a cell having only a bright cytoplasm, allowed for more accurate cell segmentation. The training set was 9 volumes consisting of three samples sets containing three 90-μm thick sub-volumes (sampling the rostral, middle, and caudal regions of the left SNc). For each sub-volume, training involved manually selecting and marking 20-30 cells with both sparse and dense distribution throughout the volume. After classifying the pixels, a quality filter of -4 and a number-of-voxels filter of 14 were applied before calculating the cell count per volume. After training, we tested the entire machine learning-based procedure on a new sample set of SNc regions expected to have an opposite cell count trend. To determine the actual cell count per SNc region, the output of the machine learning-based procedure was filtered to include only cells within the volume enclosed by the surface contours. Separate cell count values were performed for the left and right SNc regions of each brain.

**Immunohistochemistry:** Mice were transcardially perfused with 4% paraformaldehyde in 0.1 mol/L phosphate buffer. Brains were collected and post-fixed overnight in a 30% sucrose solution before being cut into 30 µm-thick sections via a cryostat. All sections were washed in PBS, incubated with 1% bovine serum albumin and 5% normal goat serum for 1 hour at room temperature. After which sections incubated with anti-sheep Tyrosine Hydroxylase (1:1,000, Novus Biologicals, Centennial, CO, NB300-110) overnight at 4°C. This was followed by incubation with either Alexa Fluor 488 goat anti-sheep IgG (1:2000, Life Technologies, A11015) secondary antibodies for 1 hour. After washing with PBS, the sections were mounted using Vectashield (Vector Laboratories, Newark, CA). Confocal image stacks were generated with an inverted A1R-HD25 confocal microscope (Nikon Instruments Inc., Melville, NY) using a 40x (NA 1.3) oil objective in 0.3 µm z-steps and at 0.43 µm/pixel.

**Neurological assessment**

Behavior assessments were performed as described previously [85-87]. Stroke-induced behavior outcomes were reported against individual pre-ischemic baseline. Analyses of behavior with the inclusion of deceased mice by assigning lowest registered scores within the cohort.

*Open field test:* To assess motor function, animals were placed in a 0.4m x 0.4m square field surrounded by a 0.4m high wall and allowed to freely explore the platform for 10 minutes [85,87]. The total distance traversed was measured, as well as central travel distance to assess anxiety-related behavior, using ANYmaze software (Stoelting Co. in Wood Dale, IL).

*Rotarod:* The mice were placed on a rotating rod (indented rod, 3.5 cm diameter) that accelerated from 4 revolutions per minute (r/min) to 80 r/min over the course of 5 minutes. The latency to fall was used to assess motor performance and was evaluated as described previously [86]. To achieve peak performance, animals were trained on the rotarod for five days with five daily trials and compared with pre-stroke baseline. The reported values are the mean of five consecutive trials.

*Grip strength:* Grip strength in the hindlimbs of the mice was measured using a grip strength meter (Columbus Instruments, Columbus, OH). The basal strength was measured the week prior to MCAO. Grip strength was assessed three times, and the average of these measurements was used as the final value.

**Statistics:** The data from the *in vivo* studies were presented using violin plots to show the distribution, mean, and interquartile range (IQR). For *in vitro* studies, the data were displayed as mean ± SD. Multigroup analyses were performed using two-way ANOVA for the effect of stroke (Contralateral vs. Ipsilateral indicated by *) and effect of conditioning (Sham conditioning vs. RIC, indicated by #), followed by *post hoc* Fisher's Least Significant Difference (LSD) test. The student's t-test was used to compare the differences between Ly6C subsets and TH+ neurons or striatal atrophy between the sham and RIC. All analyses were carried out using Prism software (GraphPad Software Inc., La Jolla, CA), and differences were considered statistically significant at $p < 0.05$.
