## Supplementary material for "Remote ischemic conditioning attenuates transneuronal degeneration and promotes stroke recovery via CD36-mediated efferocytosis": Major Resources Table are provided in the Supplemental material

In order to allow validation and replication of experiments, all essential research materials listed in the Methods should be included in the Major Resources Table below. Authors are encouraged to use public repositories for protocols, data, code, and other materials and provide persistent identifiers and/or links to repositories when available. Authors may add or delete rows as needed.

### Animals (in vivo studies)

| Species | Vendor or Source | Background Strain | Sex | Persistent ID / URL |
|---|---|---|---|---|
| C57BL/6 mouse | Jackson Laboratory | C57BL/6J | both | #000664/ www.jax.org/strain/000664 |
| CD36fl/fl mouse | Dr. Maria Febbraio, Department of Dentistry and Dental Hygiene, University of Alberta, Edmonton, Alberta, Canada | C57BL/6J | both | N/A |
| $cKO^{MM\phi}$ mouse | BNI | C57BL/6J | both | N/A |
| CD36KO mouse | Jackson Laboratory | C57BL/6J | both | #019006/ www.jax.org/strain/019006 |
| TH-GFP mouse | Dr. Kazuto Kobayashi, Department of Molecular Genetics, Institute of Biomedical Sciences, Fukushima Medical University School of Medicine, Fukushima, Japan | C57BL/6J | both | N/A |

### Genetically Modified Animals

| | Species | Vendor or Source | Background Strain | Other Information | Persistent ID / URL |
|---|---|---|---|---|---|
| **Parent - Male** | C57BL/6 mouse | Jackson Laboratory | C57BL/6J | For breeding heterozygote TH-GFP mouse line | #000664/ www.jax.org/strain/000664 |
| **Parent - Female** | TH-GFP mouse | Dr. Kazuto Kobayashi, Department of Molecular Genetics, Institute of Biomedical Sciences, Fukushima Medical University School of Medicine, Fukushima, Japan | C57BL/6J | GFP expression under the control of a modified Thy1 promoter region | N/A |

| | Species | Vendor or Source | Background Strain | Other Information | Persistent ID / URL |
|---|---|---|---|---|---|
| **Parent - Male** | $CD36^{fl/fl}$ mouse | Dr. Maria Febbraio, Department of Dentistry and Dental Hygiene, | C57BL/6J | Floxed CD36 allele | N/A |

DOI [to be added]

| | | | | | |
|---|---|---|---|---|---|
| | | University of Alberta, Edmonton, Alberta, Canada | | | |
| **Parent - Female** | LysMcre mouse | Jackson Laboratory | C57BL/6J | Cre recombinase under the LysM promoter | https://www.jax.org/strain/004781 |

**Antibodies**

| Target antigen | Vendor or Source | Catalog # | Working concentration | Lot # (preferred but not required) | Persistent ID / URL |
|---|---|---|---|---|---|
| CD45 Vioblue | Miltenyi Biotec | 130-110-664 | 1:50 | | https://www.miltenyibiotec.com/US-en/products/cd45-antibody-anti-mouse-reafinity-rea737.html#conjugate=vioblue:size=150-ug-in-1-ml |
| CD36 FITC | Miltenyi Biotec | 130-122-094 | 1:50 | | https://www.miltenyibiotec.com/US-en/products/cd36-antibody-anti-mouse-reafinity-rea1184.html#conjugate=vio-bright-b515:size=30-ug-in-200-ul |
| CD36 APC | Miltenyi Biotec | 130-122-091 | 1:50 | | https://www.miltenyibiotec.com/US-en/products/cd36-antibody-anti-mouse-reafinity-rea1184.html#conjugate=apc:size=30-ug-in-200-ul |
| CD11b PE vio770 | Miltenyi Biotec | 130-113-246 | 1:50 | | https://www.miltenyibiotec.com/US-en/products/cd11b-antibody-anti-mouse-reafinity-rea592.html#conjugate=pe-vio-770:size=150-ug-in-1-ml |
| Ly6G PE (Neutrophil) | Miltenyi Biotec | 130-123-780 | 1:50 | | https://www.miltenyibiotec.com/US-en/products/ly-6g-antibody-anti-mouse-reafinity-rea526.html#conjugate=pe:size=30-ug-in-200-ul |
| NK1.1 PE | Miltenyi Biotec | 130-120-506 | 1:50 | | https://www.miltenyibiotec.com/US-en/products/nk1-1-antibody-anti-mouse-reafinity-rea1162.html#conjugate=pe:size=30-ug-in-200-ul |
| CD45R PE (B cell) | Miltenyi Biotec | 130-110-846 | 1:50 | | https://www.miltenyibiotec.com/US-en/products/cd45r-b220-antibody-anti-mouse-reafinity-rea755.html#conjugate=pe:size=30-ug-in-200-ul |
| CD49b PE (T, NK, NKT) | Miltenyi Biotec | 130-120-969 | 1:50 | | https://www.miltenyibiotec.com/US-en/products/cd49b-antibody-anti-mouse-reafinity-rea541.html#conjugate=pe:size=30-ug-in-200-ul |

DOI [to be added]

| | | | | | |
|---|---|---|---|---|---|
| CD90.2 PE (T cell) | Miltenyi Biotec | 130-120-897 | 1:50 | | https://www.miltenyibiotec.com/US-en/products/cd90-2-antibody-anti-mouse-reafinity-rea1167.html#conjugate=pe:size=30-ug-in-200-ul |
| Ly6G APC (Neutrophil) | Miltenyi Biotec | 130-120-803 | 1:50 | | https://www.miltenyibiotec.com/US-en/products/ly-6g-antibody-anti-mouse-reafinity-rea526.html#conjugate=apc:size=30-ug-in-200-ul |
| NK1.1 APC | Miltenyi Biotec | 130-120-507 | 1:50 | | https://www.miltenyibiotec.com/US-en/products/nk1-1-antibody-anti-mouse-reafinity-rea1162.html#conjugate=apc:size=30-ug-in-200-ul |
| CD45R APC (B cell) | Miltenyi Biotec | 130-110-847 | 1:50 | | https://www.miltenyibiotec.com/US-en/products/cd45r-b220-antibody-anti-mouse-reafinity-rea755.html#conjugate=apc:size=30-ug-in-200-ul |
| CD49b APC (T, NK, NKT) | Miltenyi Biotec | 130-120-303 | 1:50 | | https://www.miltenyibiotec.com/US-en/products/cd49b-antibody-anti-mouse-reafinity-rea541.html#conjugate=apc:size=30-ug-in-200-ul |
| CD90.2 APC (T cell) | Miltenyi Biotec | 130-120-898 | 1:50 | | https://www.miltenyibiotec.com/US-en/products/cd90-2-antibody-anti-mouse-reafinity-rea1167.html#conjugate=apc:size=30-ug-in-200-ul |
| Ly6C FITC | Miltenyi Biotec | 130-111-777 | 1:50 | | https://www.miltenyibiotec.com/US-en/products/ly-6c-antibody-anti-mouse-reafinity-rea796.html#conjugate=fitc:size=150-ug-in-1-ml |
| Ly6C APC | Miltenyi Biotec | 130-111-779 | 1:50 | | https://www.miltenyibiotec.com/US-en/products/ly-6c-antibody-anti-mouse-reafinity-rea796.html#conjugate=apc:size=150-ug-in-1-ml |
| Rabbit anti-Tyrosine Hydroxylase | Thermo Fisher Scientific | 701949 | 1:1000 | | https://www.thermofisher.com/antibody/product/Tyrosine-Hydroxylase-Antibody-clone-9H8L15-Recombinant-Monoclonal/701949 |
| Sheep anti-Tyrosine Hydroxylase | Novus Biologicals | NB300-110 | 1:1000 | | https://www.novusbio.com/products/tyrosine-hydroxylase-antibody_nb300-110 |
| Mouse anti-GFP | Addgene | 180084-rAb | 1:300 | | https://www.addgene.org/180084/ |

| | | | | | |
|---|---|---|---|---|---|
| Alexa Fluor 594 donkey-anti-rabbit | Jackson ImmunoResearch | 711-587-003 | 1:50 | | https://www.jacksonimmuno.com/catalog/products/711-587-003 |
| Alexa Fluor 647 goat-anti-mouse IgG2a | Jackson ImmunoResearch | 115-607-186 | 1:250 | | https://www.jacksonimmuno.com/catalog/products/115-607-186 |
| Alexa Fluor 488 goat-anti-sheep IgG | Life Technologies | A11015 | 1:1000 | | https://www.thermofisher.com/antibody/product/Donkey-anti-Sheep-IgG-H-L-Cross-Adsorbed-Secondary-Antibody-Polyclonal/A-11015 |
| Mouse anti-CD36 | MilliporeSigma | MAB1258 | 1:2000 | | https://www.citeab.com/antibodies/225984-mab1258-anti-cd36-antibody-clone-63-mab1258 |
| Rabbit anti-GAPDH | Santa Cruz | Sc-25778 | 1:5000 | | https://www.scbt.com/p/gapdh-antibody-fl-335 |
| IRDye® 800CW Donkey anti-Mouse IgG | Licorbio | 926-32212 | 1:2000 | | https://www.licor.com/bio/reagents/irdye-800CW-donkey-anti-mouse-igg-secondary-antibody |
| IRDye® 680CW Goat anti-Rabbit IgG | Licorbio | 926-68071 | 1:2000 | | https://www.licor.com/bio/reagents/irdye-680rd-goat-anti-rabbit-igg-secondary-antibody |

### DNA/cDNA Clones

| Clone Name | Sequence | Source / Repository | Persistent ID / URL |
|---|---|---|---|
| N/A | N/A | N/A | N/A |

### Cultured Cells

| Name | Vendor or Source | Sex (F, M, or unknown) | Persistent ID / URL |
|---|---|---|---|
| Splenocyte; primary culture | C57BL/6 mouse | M, F | |
| Splenocyte; primary culture | cKO$^{MM\phi}$ mouse | M, F | |
| Brain immune cell | | | |

### Data & Code Availability

| Description | Source / Repository | Persistent ID / URL |
|---|---|---|
| RNA-seq data | National Center for Biotechnology Information Short Read Archive | accession number: PRJNA525413 |

### Other

| Description | Source / Repository | Persistent ID / URL |
|---|---|---|
| CD11b Microbeads | Miltenyi Biotec | https://www.miltenyibiotec.com/US-en/products/cd11b-microbeads-human-and-mouse.html#130-049-601 |
| MS Columns | Miltenyi Biotec | https://www.miltenyibiotec.com/US-en/products/ms-columns.html#130-042-201 |

DOI [to be added]

| | | |
|---|---|---|
| Nueral Tissue Dissociation Kit | Miltenyi Biotec | https://www.miltenyibiotec.com/US-en/products/neural-tissue-dissociation-kits.html#130-092-628 |
| Debris removal kit | Miltenyi Biotec | https://www.miltenyibiotec.com/US-en/products/debris-removal-solution.html#130-109-398 |
| PKH67 Green Fluorescent Cell Linker | Sigma-aldrich | https://www.sigmaaldrich.com/US/en/product/sigma/mini67?srsltid=AfmBOoqtpZB2j882TS6U_WhFZMhuDBPieZkKADVJ_B-U_tMT_ruN2CME |
| FluoSpheres™ Polystyrene Microspheres, 1.0 µm, red fluorescent (580/605) | Thermo Fisher | https://www.thermofisher.com/order/catalog/product/F13083 |

DOI [to be added]
